## Supplementary Information for "METASPACE-ML: Context-specific metabolite annotation for imaging mass spectrometry using machine learning"

### Supplementary Note 1: Reprocessing training datasets with the same set of METASPACE parameters

Following the selection of training datasets from the METASPACE knowledge base, each dataset was reprocessed using the rule-based MSM approach with the following modifications to the preset configuration file:

1. In positive and negative polarity modes, we have considered the following target adducts, respectively : ('+H' , +Na', '+K') and ('-H', '+Cl').
2. The decoy sample size of 20
3. m/z tolerance of 3 ppm

### Supplementary Note 2: Construction of the CoreMetabolome database

The CoreMetabolome database was generated by first downloading the HMDB v4 database (Wishart et al. 2018) as a comprehensive source of data. Other databases were used to "white-list" HMDB_v4 molecules into the CoreMetabolome databases as follows:

1. The subset of HMDB_v4 which was tagged as either "Detected" or "Quantified" was added into the CoreMetabolome database, under the assumption that previously observed molecules will likely be observed again.
2. The following resources were downloaded:
    a. HMDB (Wishart et al. 2018), ECMDB (Sajed et al. 2016) , YMDB (Ramirez-Gaona et al. 2017) : Human, *E. coli*, and yeast. Any conserved molecules across all three divergent species were deemed "core" and added to the CoreMetabolome_v1 database.
    b. KEGG (data/kegg_bio_cmpds.json). KEGG is a well-curated manually annotated database of biochemical pathways including compounds.
    c. LipidMaps, a moderate-sized, well-annotated and curated lipids database was downloaded on (July 13th 2022).
    d. MSMLS, a large collection of commercially available metabolite standards. The commercial availability for standards is a good proxy for biological importance and ubiquitous presence in biology.
    e. An in-house library of commercially available metabolite standards within the Alexandrov laboratory.
    f. Mayo clinic targeted metabolites were included under the assumption that metabolites run for clinically relevant medical sets are likely important.
3. Together, the resources in step 2 were preprocessed and combined to create the **CoreMetabolome _*v1*** database. For reproducibility, please check the following jupyter notebook (https://github.com/DinosaurInSpace/core_metabolome/blob/master/core_metabolome_reproduce.ipynb) .

4. When testing the **CoreMetabolome _v1** database, it was noticed some formulas failed, as they were present in the database as charged compounds, not neutrals and it was corrected accordingly. In addition, mono- and divalent metal cations (eg. $Ca^{2+}$) with a low likelihood of direct detection in mass spectrometry experiments were also dropped. The database without these ions formed the **CoreMetabolome_v2** database.
5. The **CoreMetabolome_v2** database was then manually reviewed based on CMR's subject matter expertise, and the free-text metadata in HMDB.
6. Molecules which were not obviously of biological origin were dropped including:
    a. Drugs and drug metabolites. Frequently described as "...observed in patients treated with…."
    b. Dietary molecules only present in food. Frequently described as "...may be a biomarker for…."
    c. Hypothetical molecules. Frequently described as "...predicted by biotransformer…."
    d. Molecules without any descriptive information linking them to biological context.
7. The remaining manual-reviewed molecules formed the **CoreMetabolome_v3** database.
8. The **CoreMetabolome_v3** [11,440 metabolites], is available here: https://github.com/DinosaurInSpace/core_metabolome/blob/master/coremetabolome_curated/core_metabolome_v3.csv

9. Note: Exclusion of particular metabolites from ***CoreMetabolome_v3*** is made without negative judgement, and instead highlights some of the challenges of database conception versus use. For example, if looking for novel metabolites it is useful to have comprehensive databases of every molecule that is known to or reasonably predicted to exist. However, in target-decoy approaches the target database should be most reflecting the possible metabolome of a sample whereas including infeasible molecules (e.g. exogenous molecules where they are not expected) can lead to incorrect estimation of FDR. This phenomenon is well-known in the field of proteomics, where searching against sample-specific databases is the norm.

### Supplementary Note 3: Calculation of ion coverage in testing datasets

To assess the coverage of target ions in each testing dataset based on their presence in any of the training datasets, we developed a straightforward measure. An ion in the testing dataset was considered covered if it was found in any of the training datasets. We then determined the coverage for each testing dataset by calculating the proportion of ions that were covered.

Specifically, for each ion *i* within a given dataset *ds*, we determined whether it was covered by the training data ($n_{dss} > 0$). We defined an indicator function *C(i)* which equals 1 if ion *i* was covered and 0 otherwise.

The coverage for each testing dataset *ds* was then calculated using the following formula:

$$C(i) = \left\{ 1\ if n_{dss} > 0,\ 0\ otherwise \right\}$$

$$coverage(ds) = \frac{1}{N(ds)} \sum_{i \in I(ds)} C(i)$$

where *N(ds)* represents the total number of unique target ions for testing dataset *ds*, *I(ds)* is the set of target ions for that dataset, and $n_{dss}$ is the number of training datasets where ion *i* has been observed.

The coverage value ranges from 0 to 1, where 0 indicates that none of the ions are covered and 1 indicates that all ions are covered. The coverage scores for each testing dataset is provided in **Supplementary Data 5.**

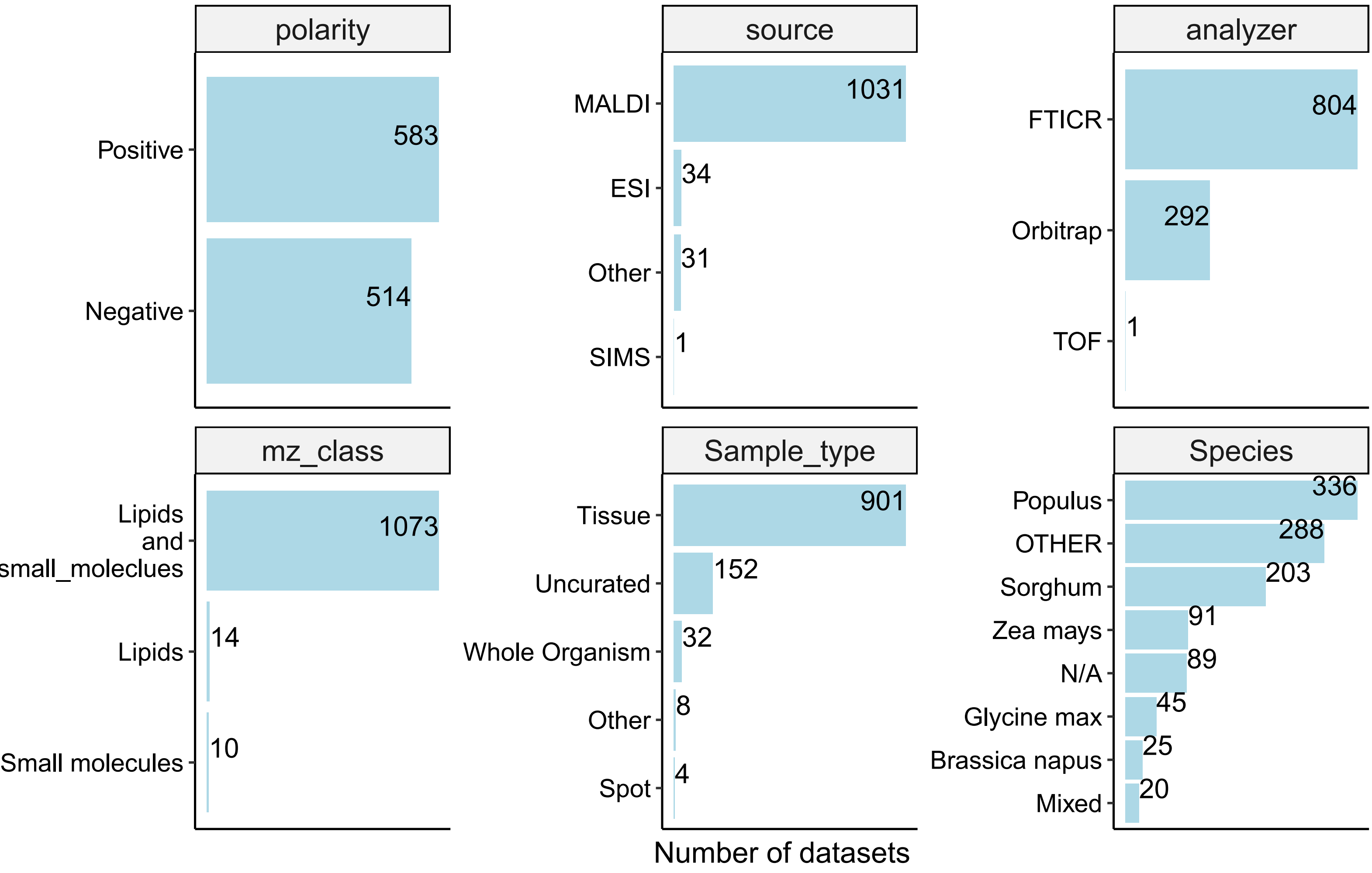


***Supplementary Fig. 1: Breakdown of METASPACE plant datasets by context***

Breakdown of the pool of plant datasets by their associated technical and biological metadata making up the contexts (see methods). Number of datasets are shown on the x-axis and the classes in each context are shown on the y-axis

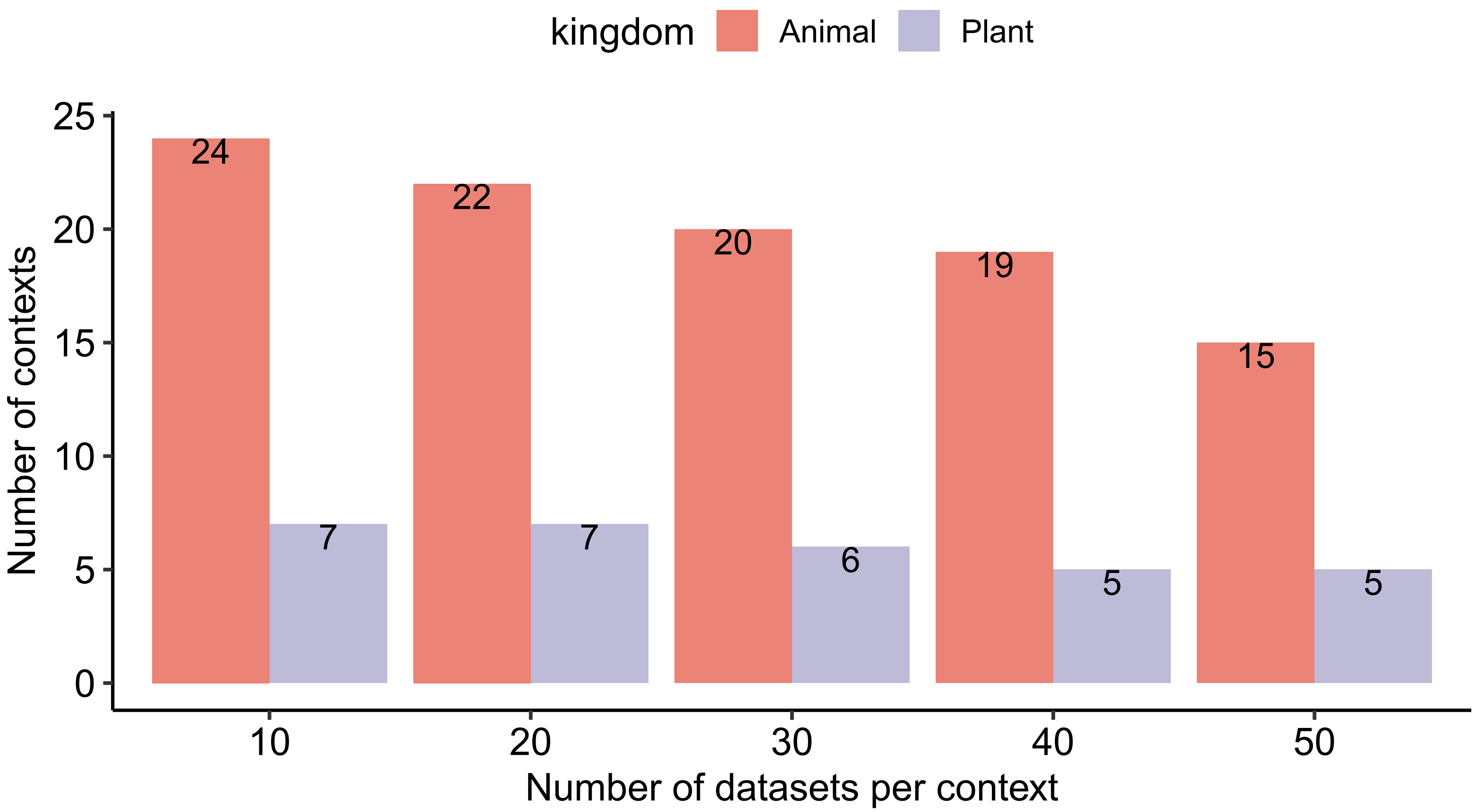


***Supplementary Fig. 2: Number of training datasets per context and kingdom***

Bar plot showing the number of training datasets per context in 5 different models with varying sizes (number of datasets per context. The number of datasets per context range from 10-50 are displayed on the x-axis and the number of different contexts represented in each set is shown on the y-axis. Colors represent the kingdom (either animal or plant). The number of contexts are displayed on top of each bar.

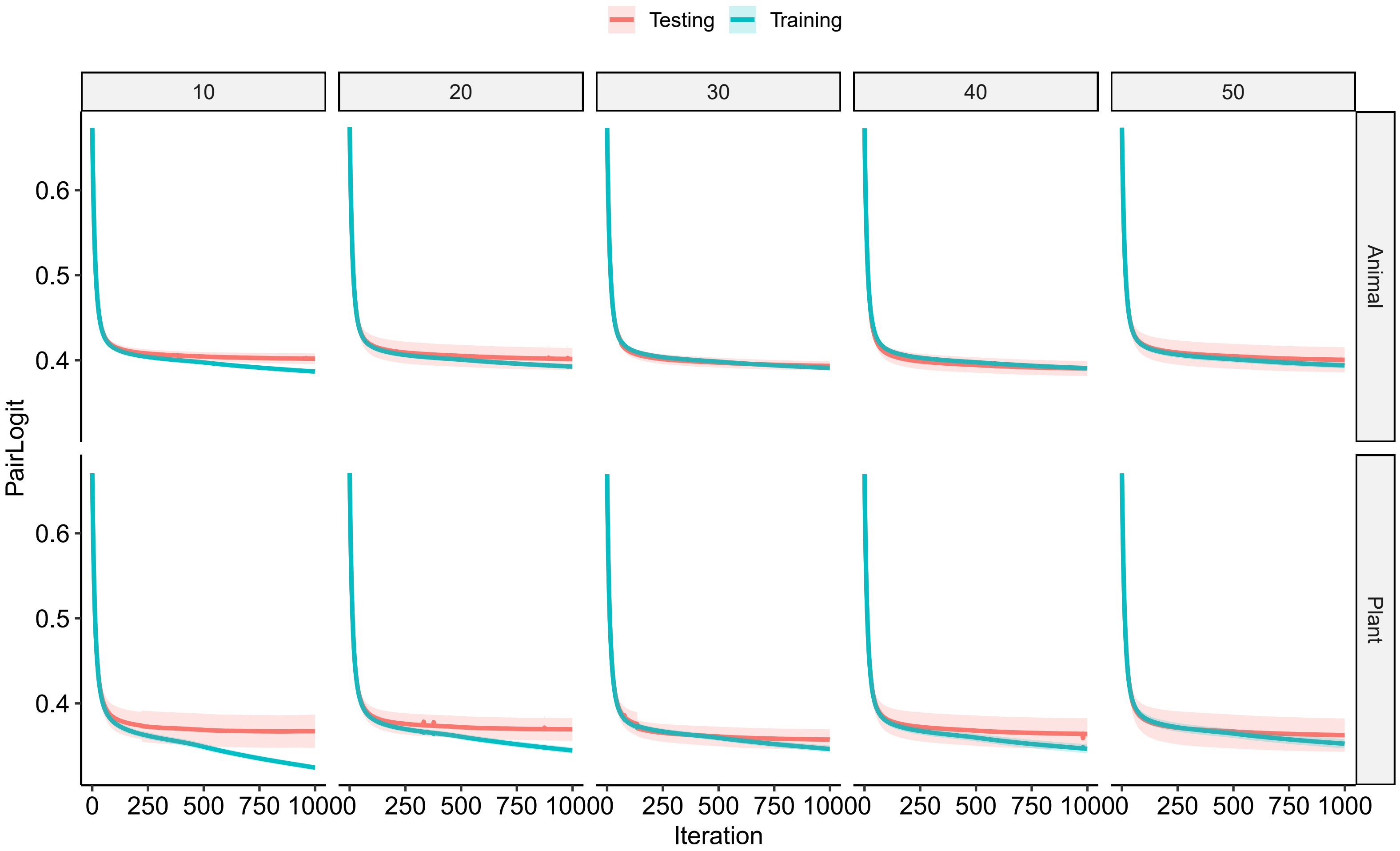


***Supplementary Fig. 3: Model learning curves***

Line plot showing how the PairLogit error decreases for each of the 1000 training iterations. Columns show the number of datasets per context (see Supplementary Fig. 2) for 5 different models. Rows show results for each kingdom (either animal or plant). Training and cross-validation data are colored blue and red, respectively. Shades around the curve represent standard deviation across 5 different cross validation splits (see methods).

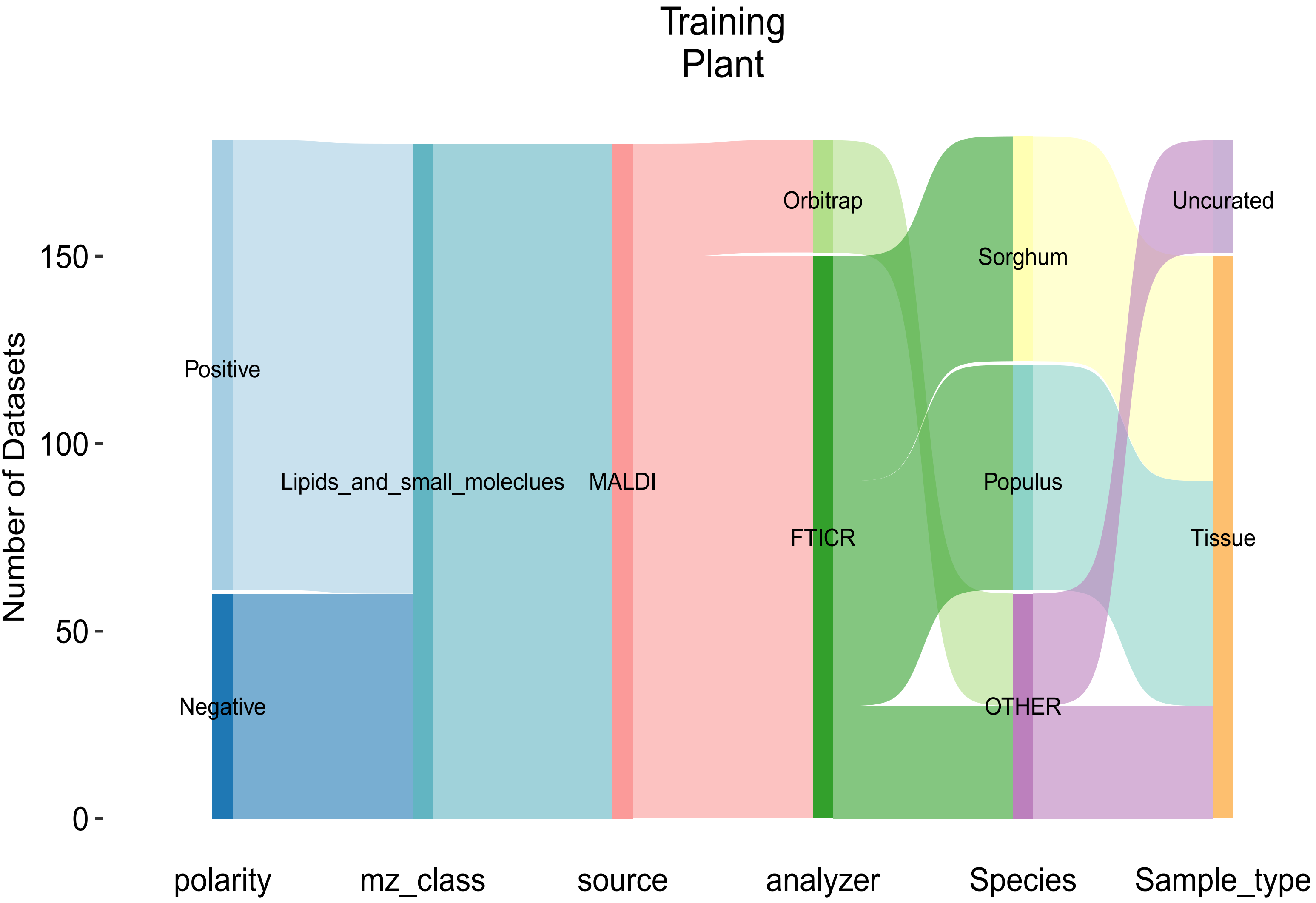


***Supplementary Fig. 4: Breakdown of plant training datasets***

Sankey diagrams showing the breakdown of training plant-based datasets by different parameters. Properties of the datasets are described on the x-axis and the corresponding number of datasets in each level is plotted on the y-axis. Each property can be divided into multiple classes which are represented by the colors of the nodes and their corresponding flow to the next layer. A flow from the first to the last node represents a single context

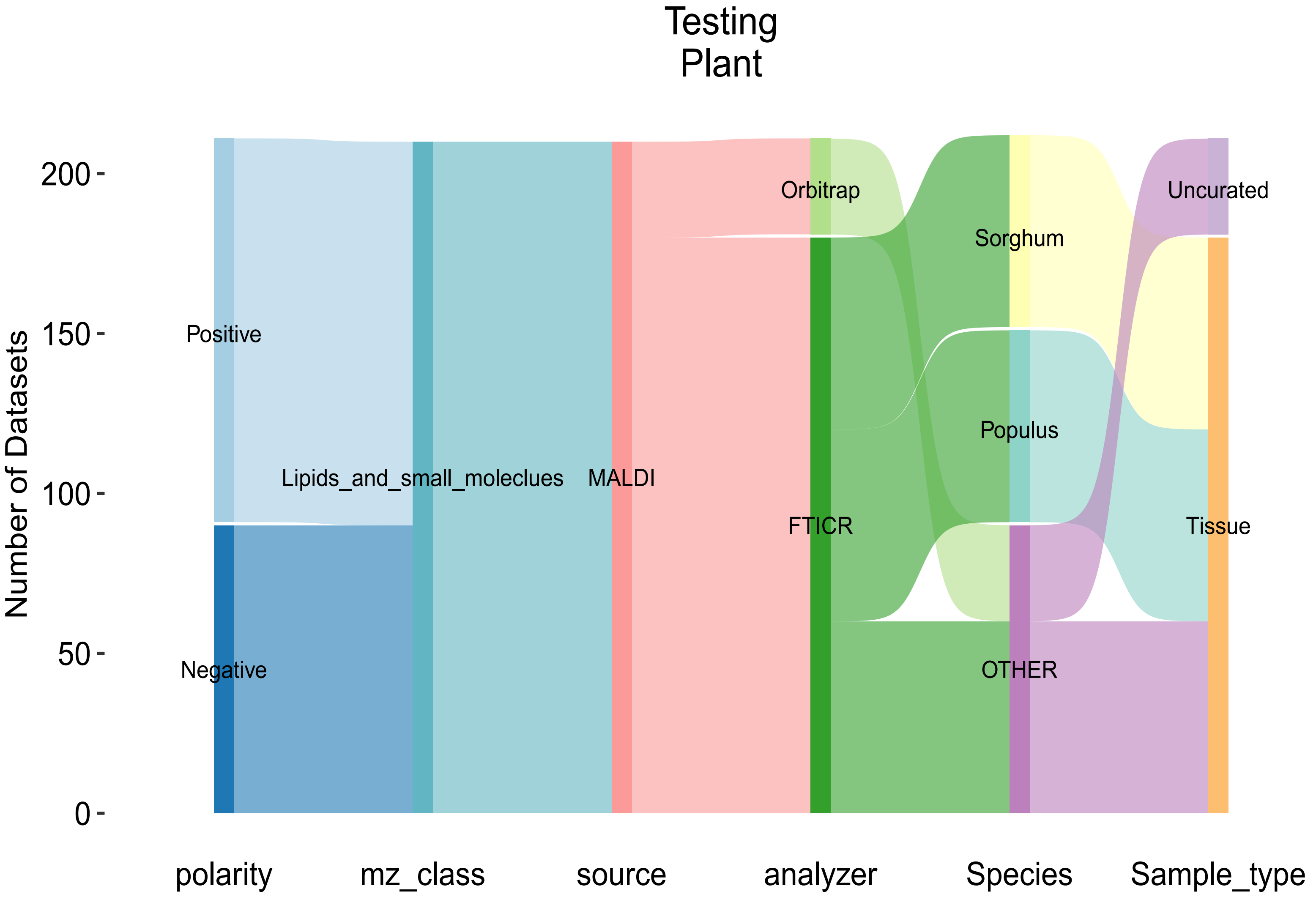


***Supplementary Fig. 5: Breakdown of plant testing datasets***

Sankey diagrams showing the breakdown of testing plant-based datasets by different parameters. Properties of the datasets are described on the x-axis and the corresponding number of datasets in each level is plotted on the y-axis. Each property can be divided into multiple classes which are represented by the colors of the nodes and their corresponding flow to the next layer. A flow from the first to the last node represents a single context

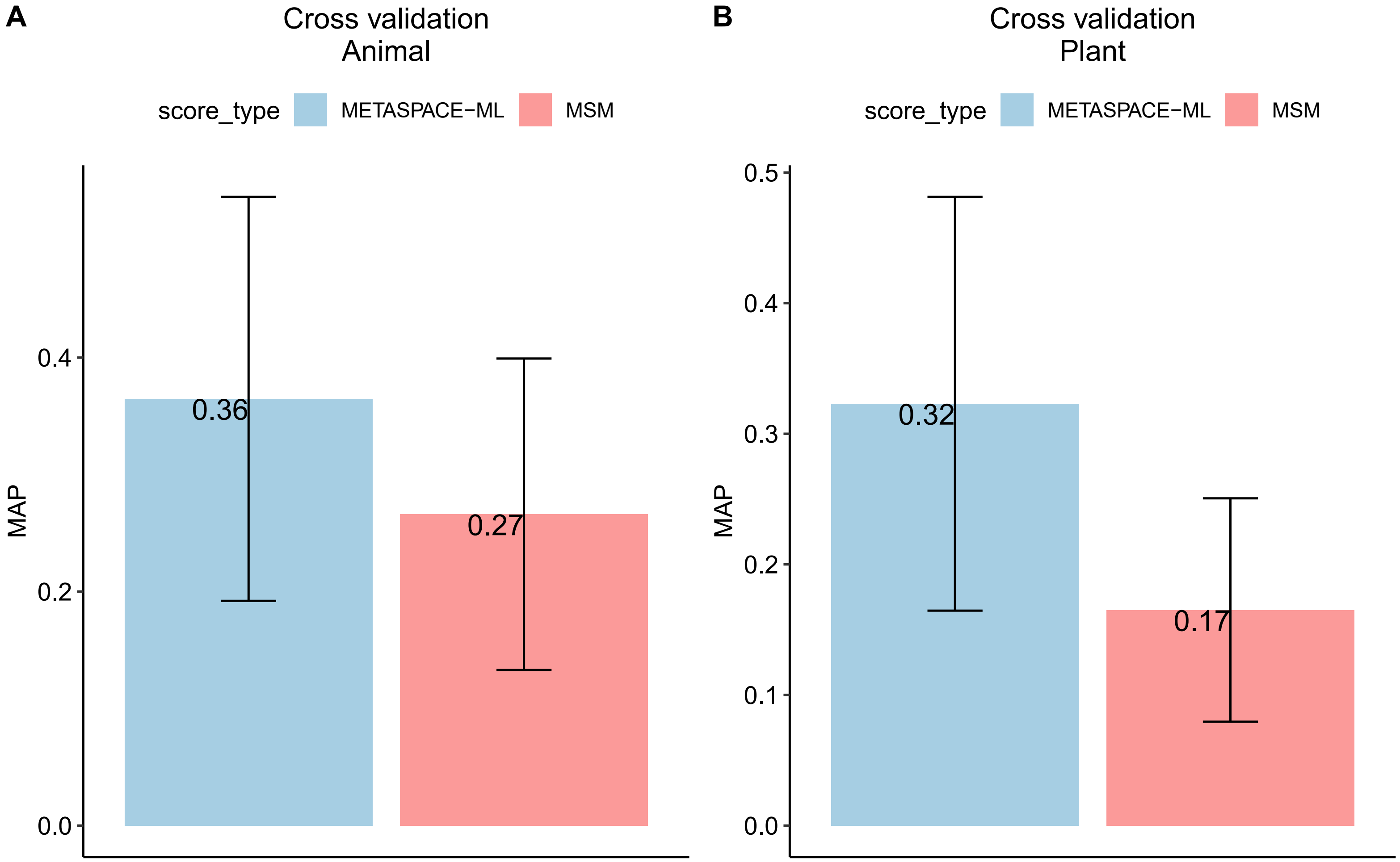


***Supplementary Fig. 6: MAP scores for cross validation per kingdom***

(A) and (B) Bar graph showing MAP scores for cross validated datasets for animal (A) and plant (B) datasets. Bars are colored based on the approach type. AP (average precision) scores are calculated for each group (dataset + adduct) and the score is the average across all groups for a given database. Error bars represent +- standard deviation of AP scores. MAP scores are displayed on the top of each bar.

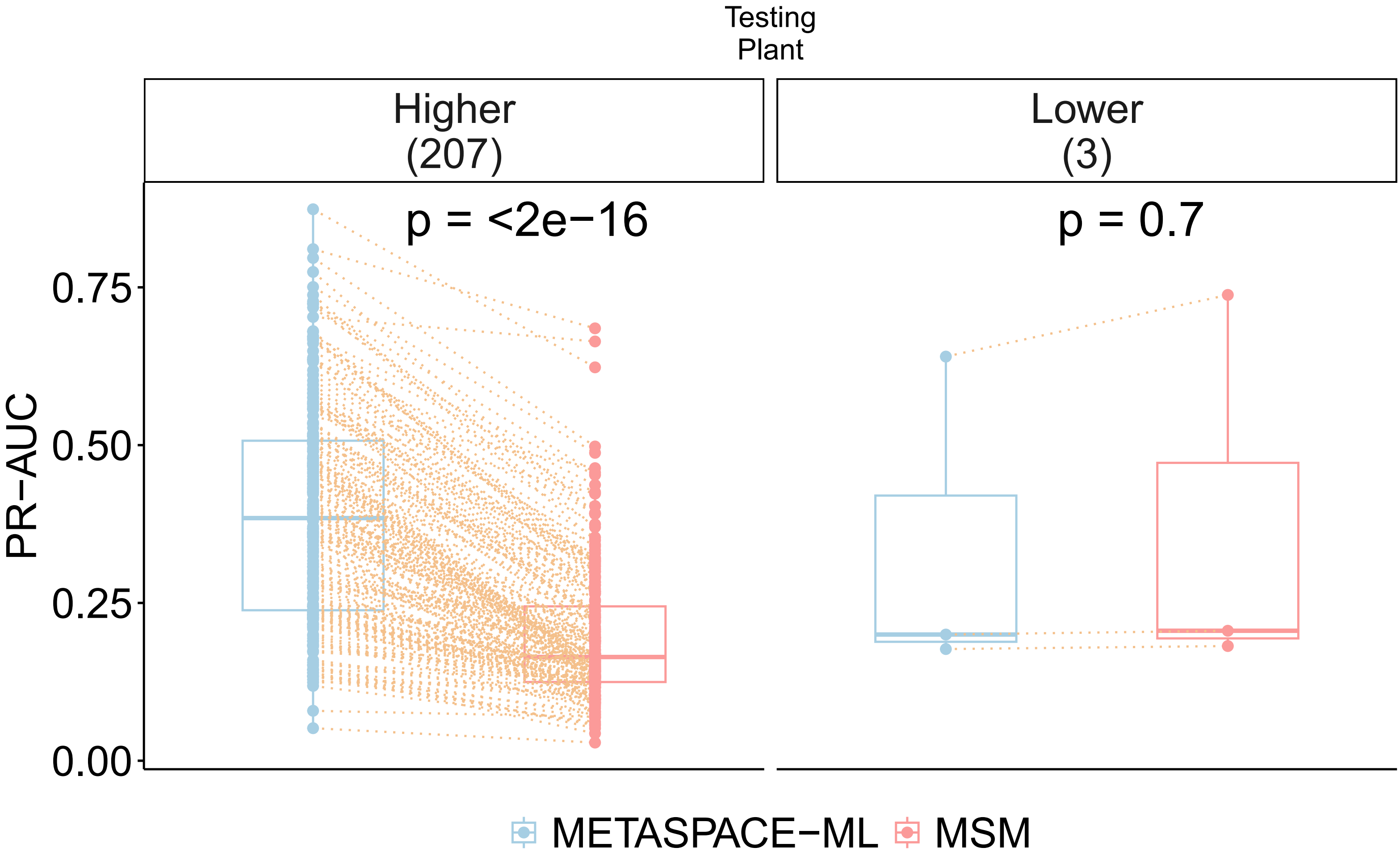


***Supplementary Fig. 7: Distribution of area under precision-recall curve for plant testing datasets***

Paired boxplot showing area under precision-recall curve (AUC) per dataset for plant testing datasets. X-axis represents the approach type and the y-axis represents AUC scores. Each dot represents a dataset and an orange dotted edge is drawn between the same datasets from both approaches. Boxplot are grouped by the difference of AUC scores (Delta) where positive differences denote higher AUC in Metaspace-ML relative to MSM and vice-versa. The number of datasets in each group are shown in parentheses. Exact p-values are based on a two-tailed paired Wilcoxon signed-rank test between AUC scores across both approaches. The boxplots' bottom and top edges represent the 25th and 75th percentiles, with the median (50th percentile) line inside the box. Whiskers are omitted; minimum and maximum values are represented by jittered data points.

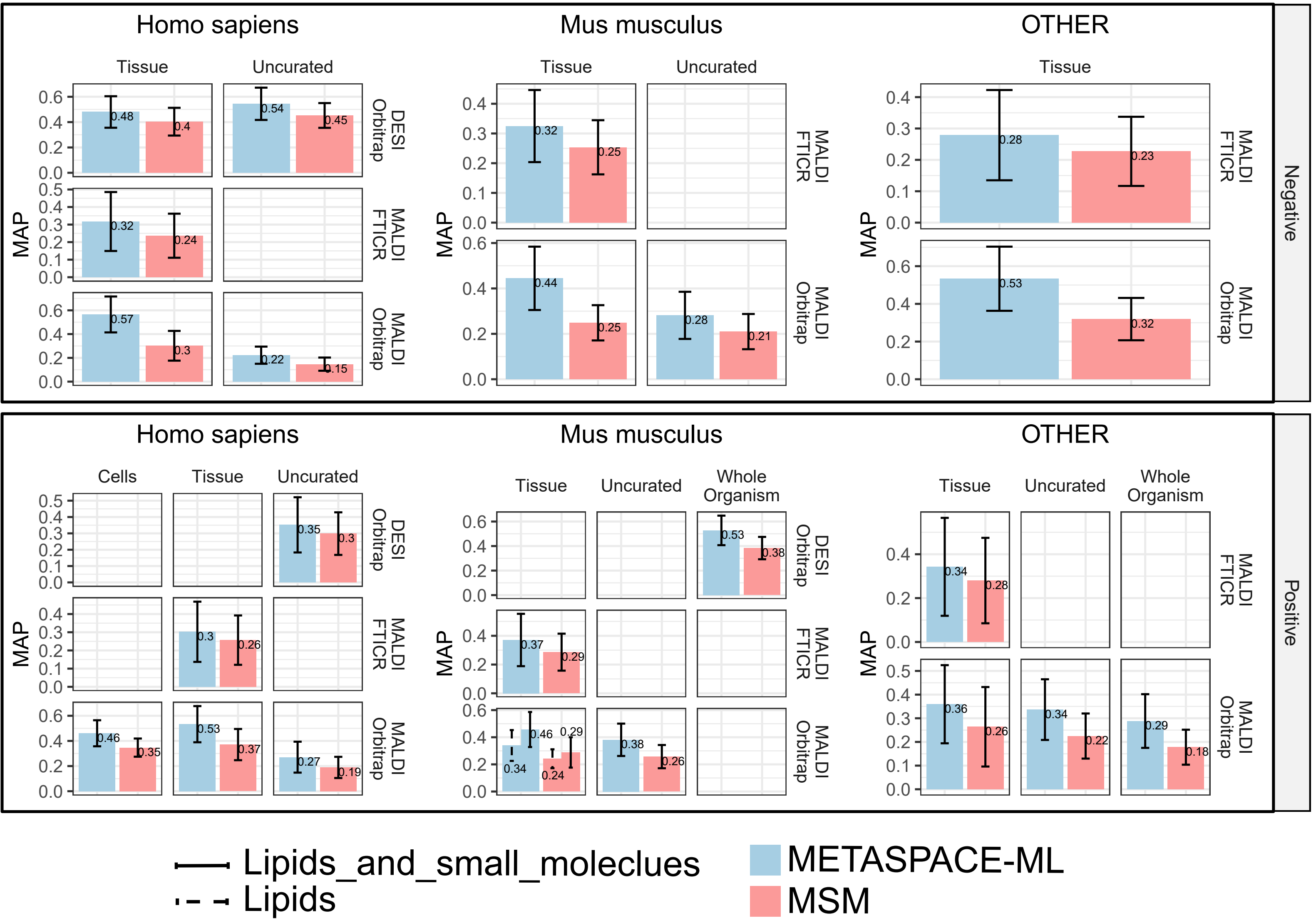


***Supplementary Fig. 8: Context-specific MAP scores for animal testing datasets***

Each bar graph shows MAP scores for 30 datasets representing a given context. Upper and lower panels represent negative and positive polarity, respectively. Each panel is divided into 3 sub panels based on the organism and each subpanel is a grid where columns and rows represent sample type and ionization source + analyzer, respectively. Bars are colored based on the approach type. AP (average precision) scores are calculated for each group (dataset + adduct) and the score is the average across all groups for a given database. Error bars represent +- standard deviation of AP scores. MAP scores are displayed on the top of each bar.

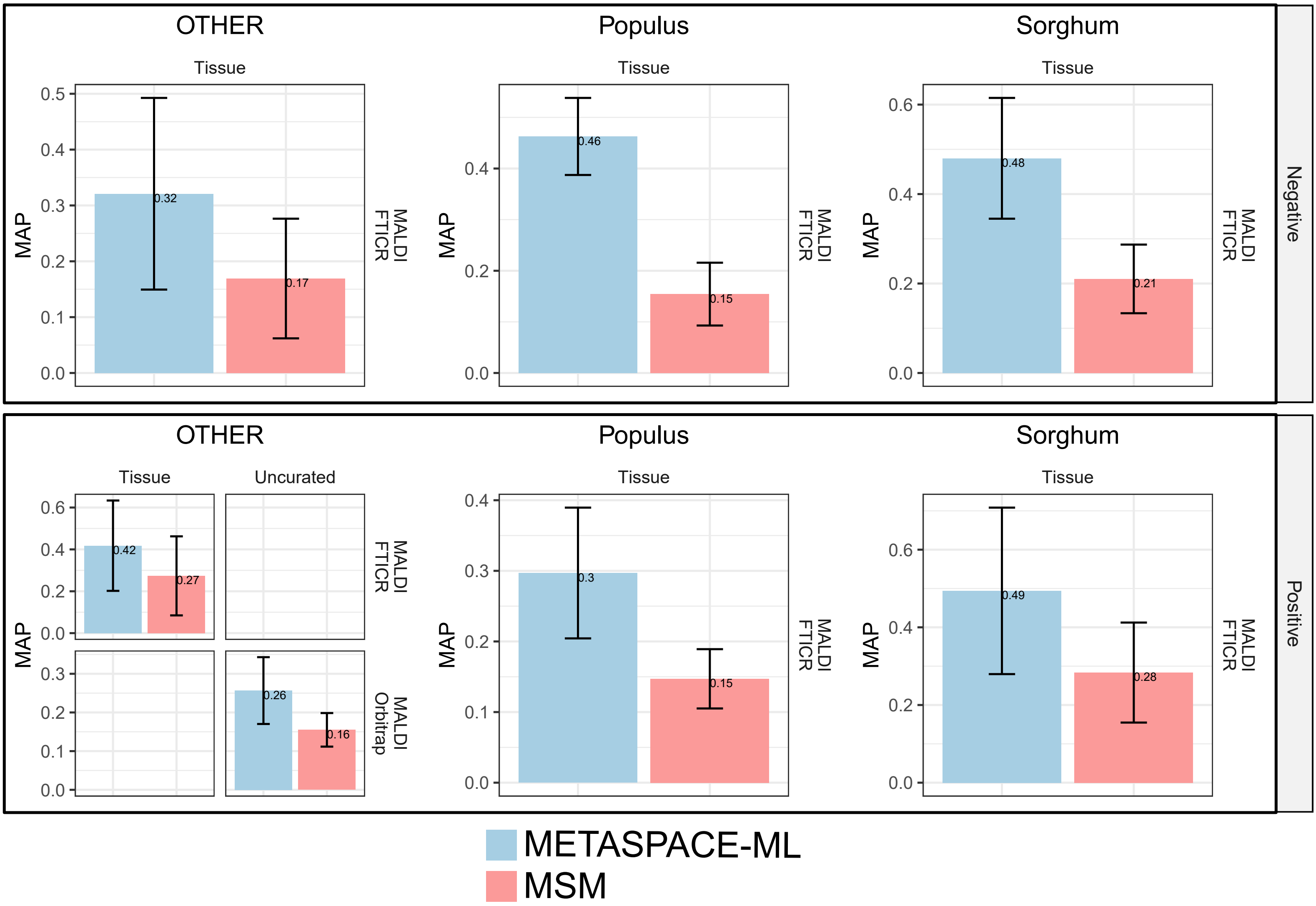


***Supplementary Fig. 9: Context-specific MAP scores for plant testing datasets***

Each bar graph shows MAP scores for 30 datasets representing a given context. Upper and lower panels represent negative and positive polarity, respectively. Each panel is divided into 3 sub panels based on the organism and each subpanel is a grid where columns and rows represent sample type and ionization source + analyzer, respectively. Bars are colored based on the approach type. AP (average precision) scores are calculated for each group (dataset + adduct) and the score is the average across all groups for a given database. Error bars represent +- standard deviation of AP scores. MAP scores are displayed on the top of each bar.

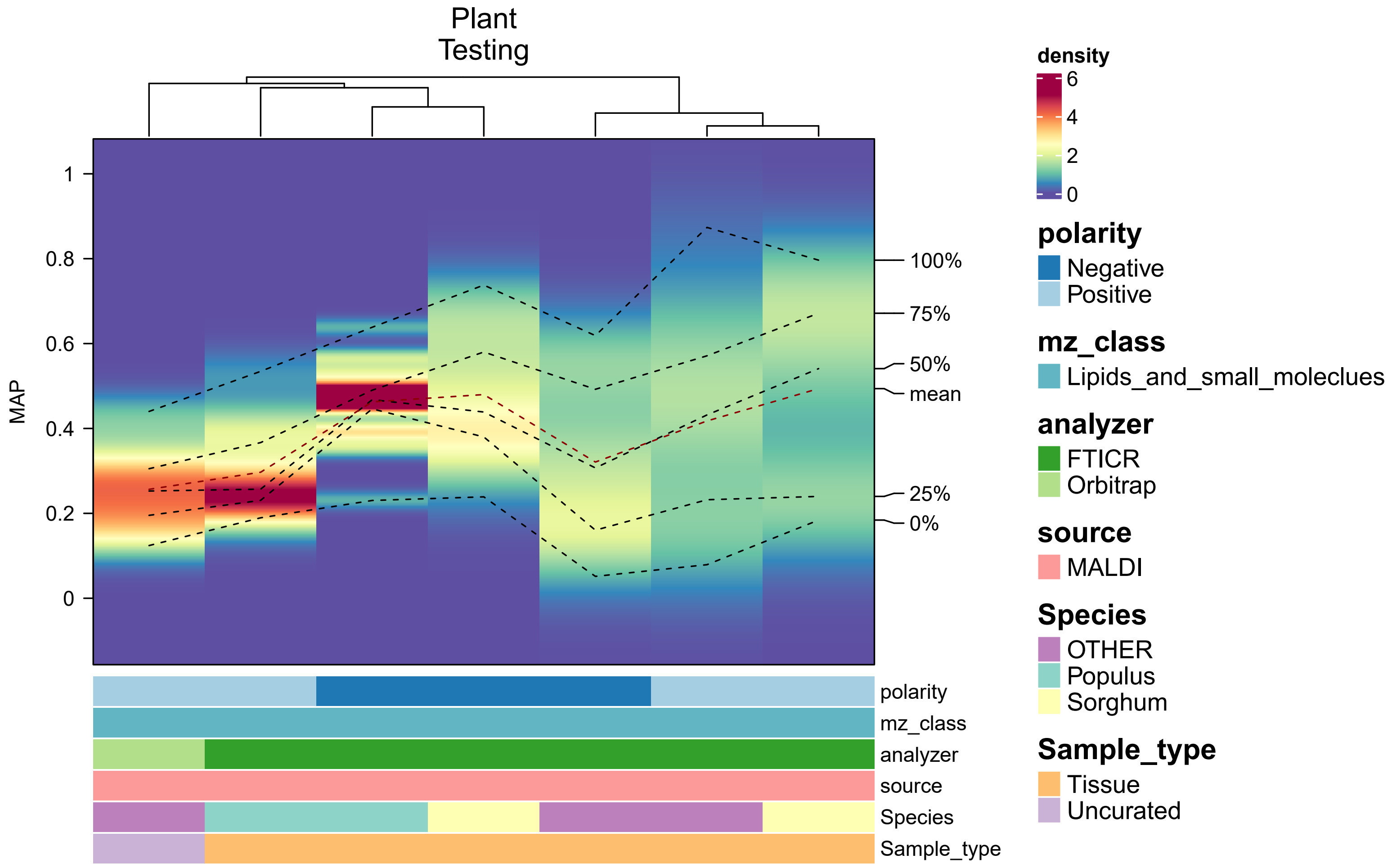


***Supplementary Fig. 10: Context-specific distribution of MAP scores for plant testing datasets***

Density heatmap showing the distribution of MAP scores across datasets for each context in plant-based testing datasets. Each column represents a context which is described by its constituent metadata as 1D annotation bars colored by the classes in each metadata variable. The y-axis shows the MAP scores and the color gradient represents their density. Columns are hierarchically clustered using a distance metric based on the Kolmogorov-Smirnov statistic. Dotted lines in each column represent different quantiles of the MAP scores in addition to the mean.

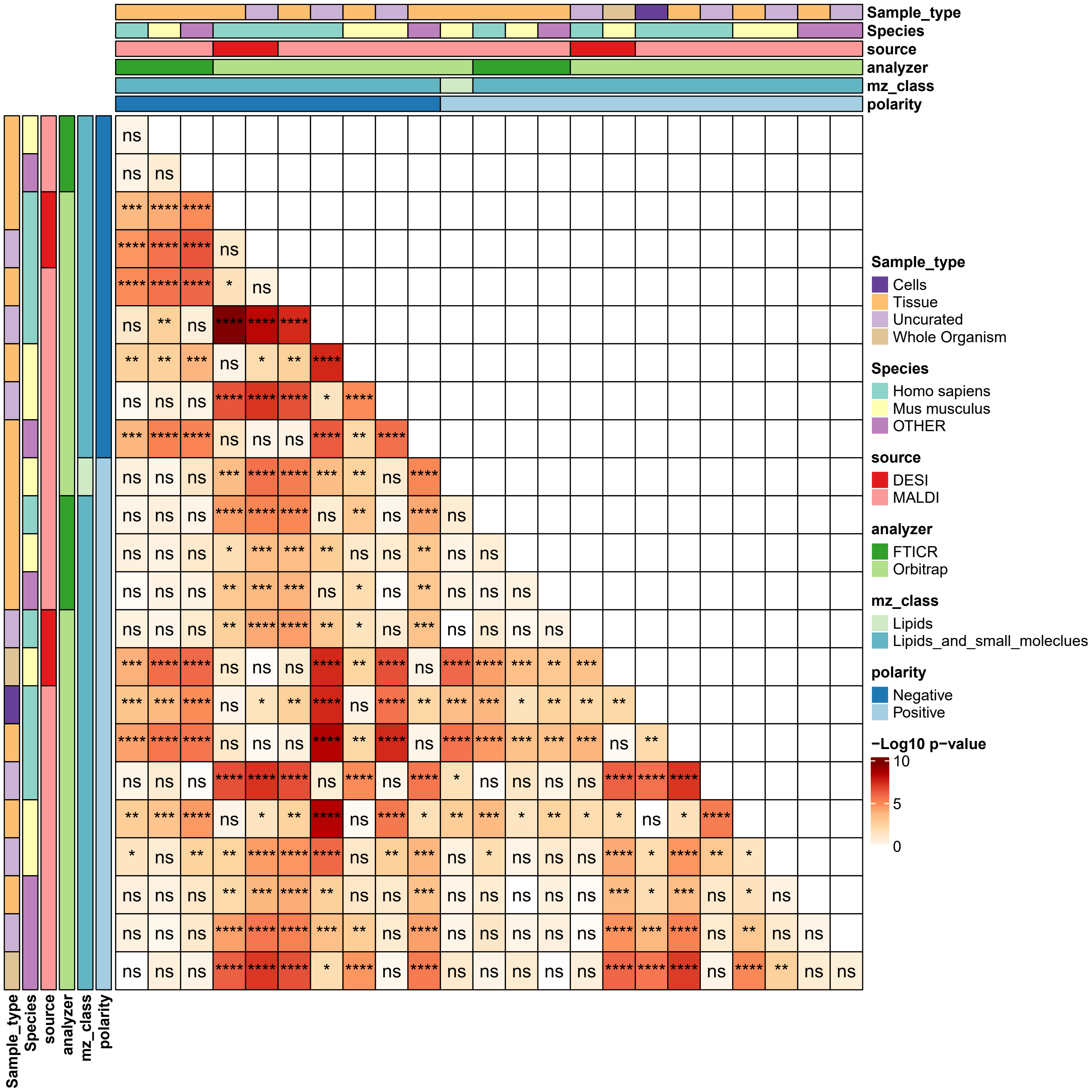


***Supplementary Fig. 11: Comparison of MAP scores between contexts in animal testing datasets***

Asymmetric heatmap showing the significance analysis (using Wilcoxon test) results for MAP scores comparison between each pair of contexts in animal testing datasets. Each column and row represents a context which is described by its constituent metadata as 1D annotation bars colored by the classes in each metadata variable. Color gradient corresponds to -Log10 p-value. Asterisks are added to denote significance level (**** < 0.0001, *** < 0.001, ** < 0.01, * < 0.05, ns >= 0.05) based on the Wilcoxon test.

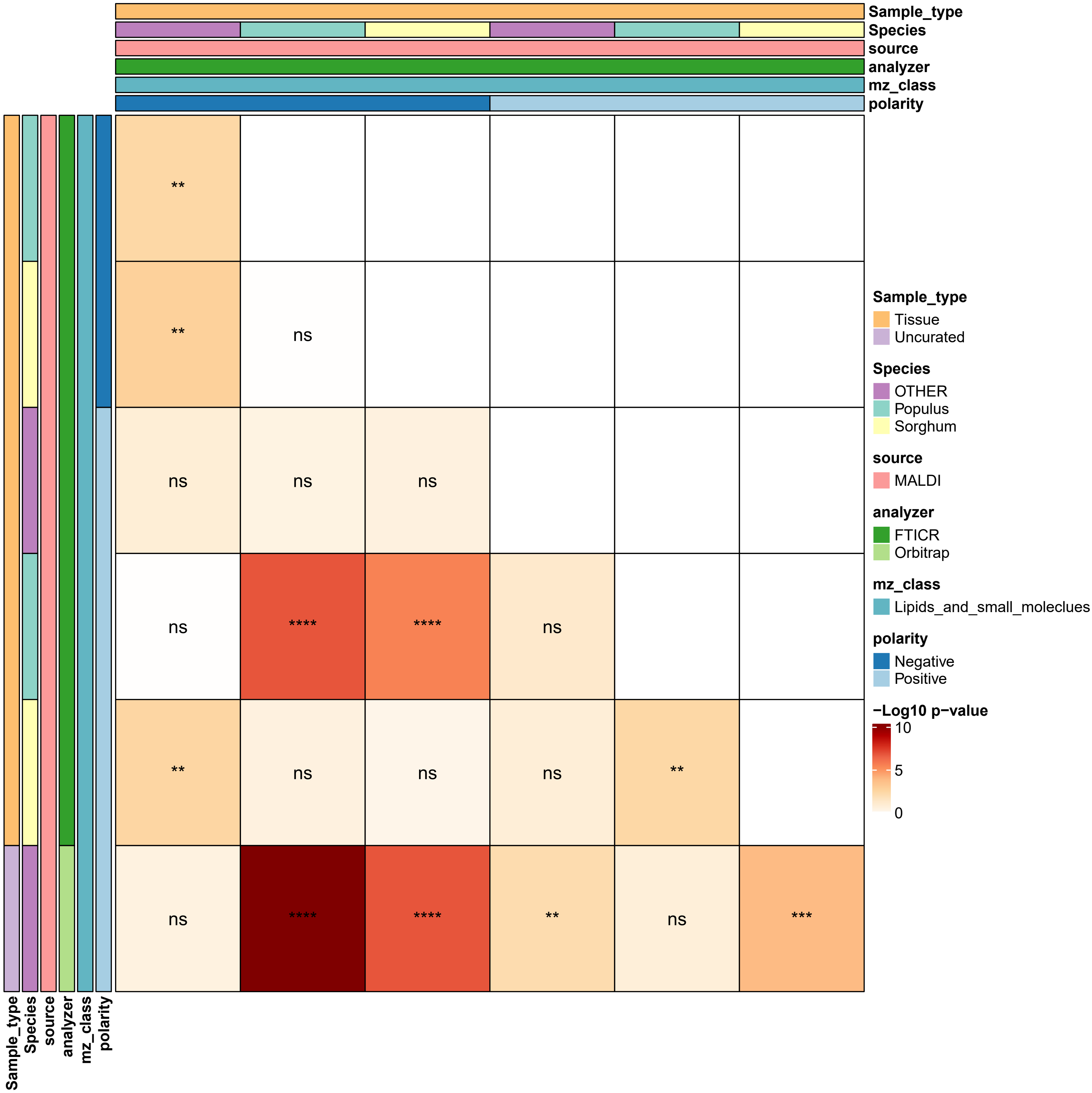


***Supplementary Fig. 12: Comparison of MAP scores between contexts in plant testing datasets***

Asymmetric heatmap showing the significance analysis (using Wilcoxon test) results for MAP scores comparison between each pair of contexts in plant testing datasets. Each column and row represents a context which is described by its constituent metadata as 1D annotation bars colored by the classes in each metadata variable. Color gradient corresponds to -Log10 p-value. Asterisks are added to denote significance level (**** < 0.0001, *** < 0.001, ** < 0.01, * < 0.05, ns >= 0.05) based on the Wilcoxon test.

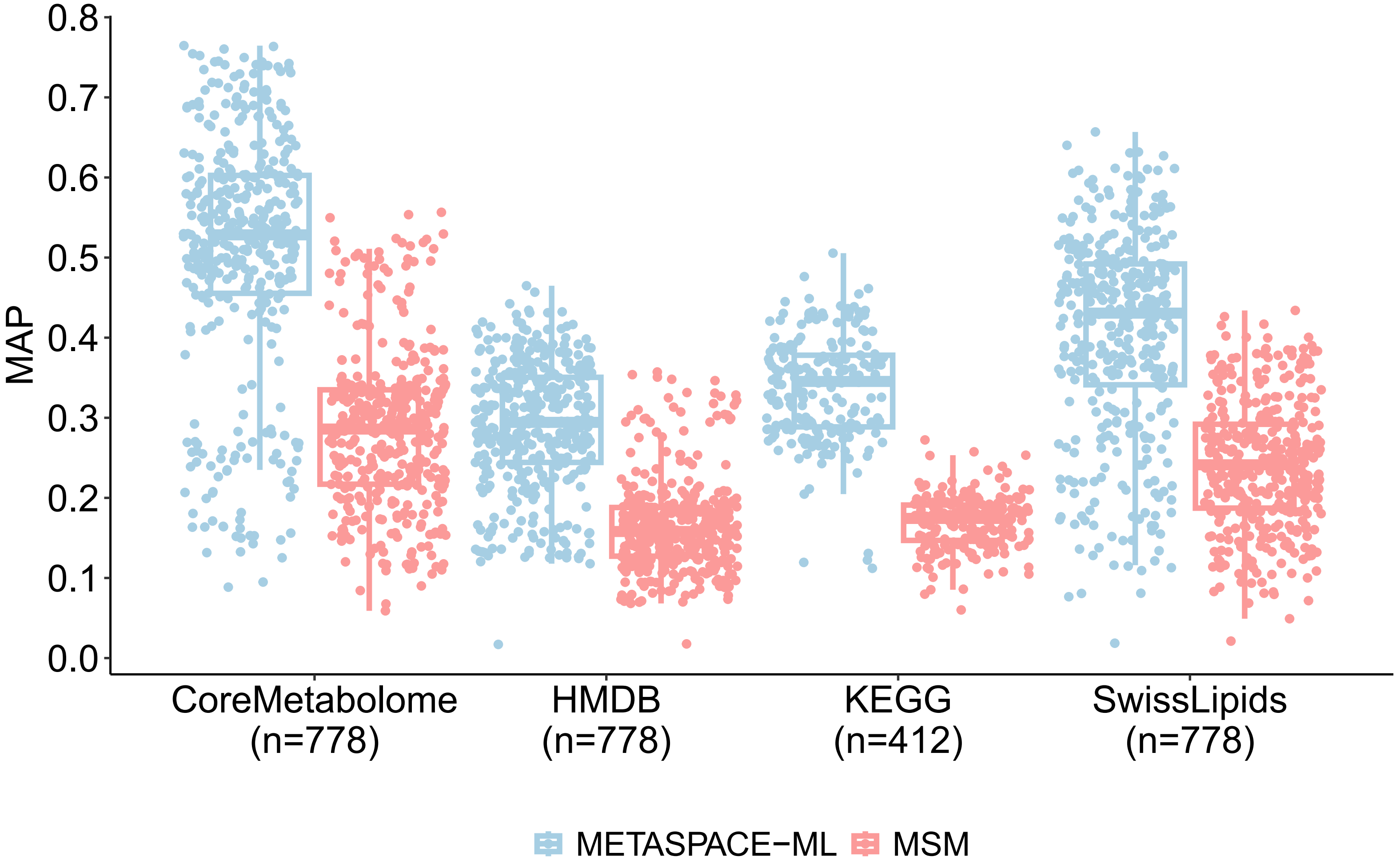


***Supplementary Fig. 13: Distribution of MAP scores in animal testing datasets per annotation database***

Boxplot showing MAP scores across different annotation databases. Each dot is a dataset and color corresponds to either METASPACE-ML or MSM-based approach. Number of datasets are displayed below x-axis labels. Boxplots' bottom and top edges represent the 25th and 75th percentiles, with the median (50th percentile) line inside the box. Whiskers extend to the minimum and maximum values within 1.5 times the interquartile range from the quartiles; the minimum and maximum values are represented by the extent of the jittered data points.

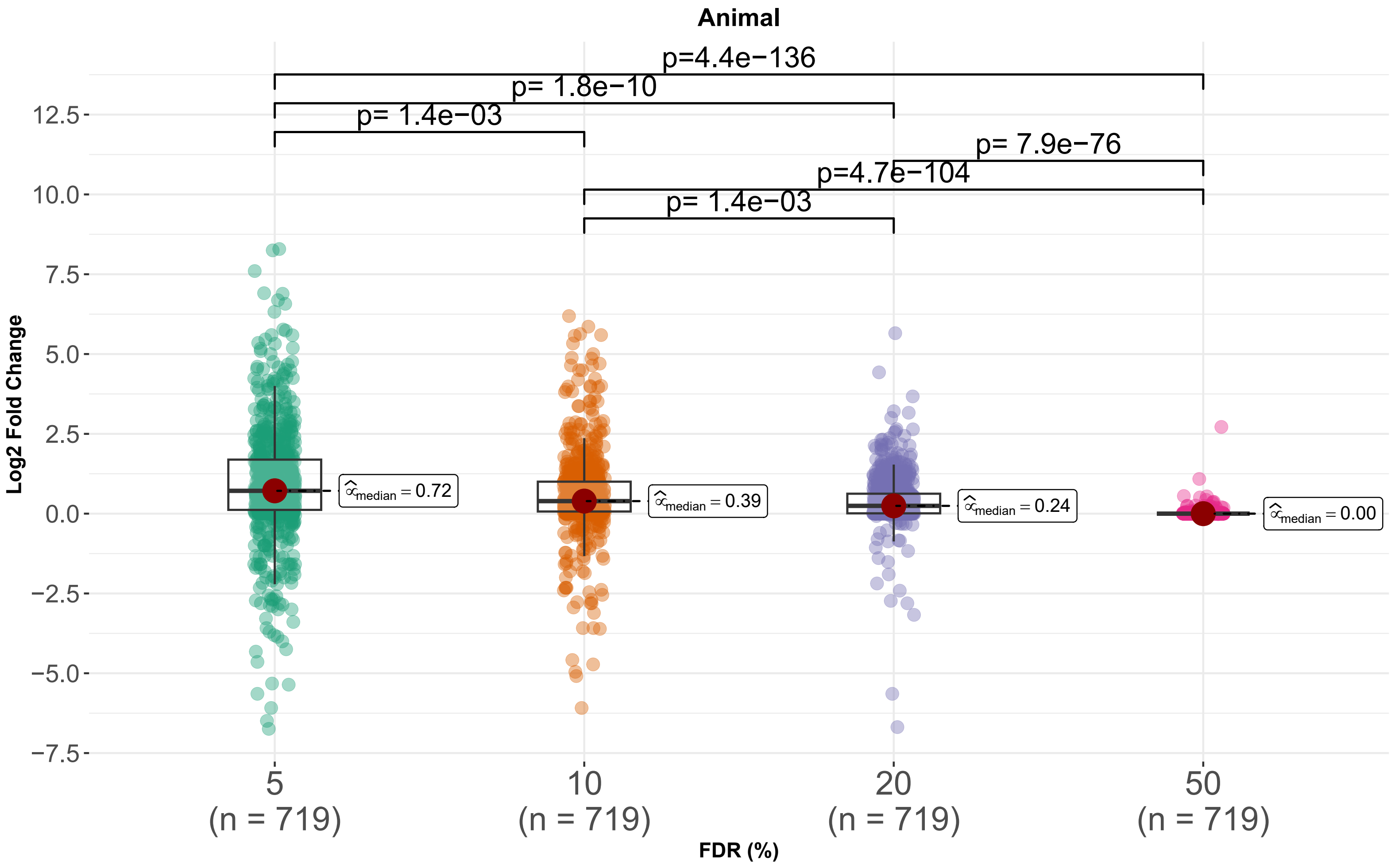


***Supplementary Fig. 14: Distribution of LFC in animal testing datasets across FDR thresholds***

Box plot showing distribution of Log2 fold changes of the number of annotations of METASPACE-ML relative to rule-based approach across all animal testing datasets, for different FDR thresholds. A dot corresponds to a test dataset. Exact p-values from a two-tailed Wilcoxon rank-sum test are shown above each comparison. Boxplots' bottom and top edges represent the 25th and 75th percentiles, with the median (50th percentile) line inside the box. Whiskers extend to the minimum and maximum values within 1.5 times the interquartile range from the quartiles; the minimum and maximum values are represented by the extent of the jittered data points.

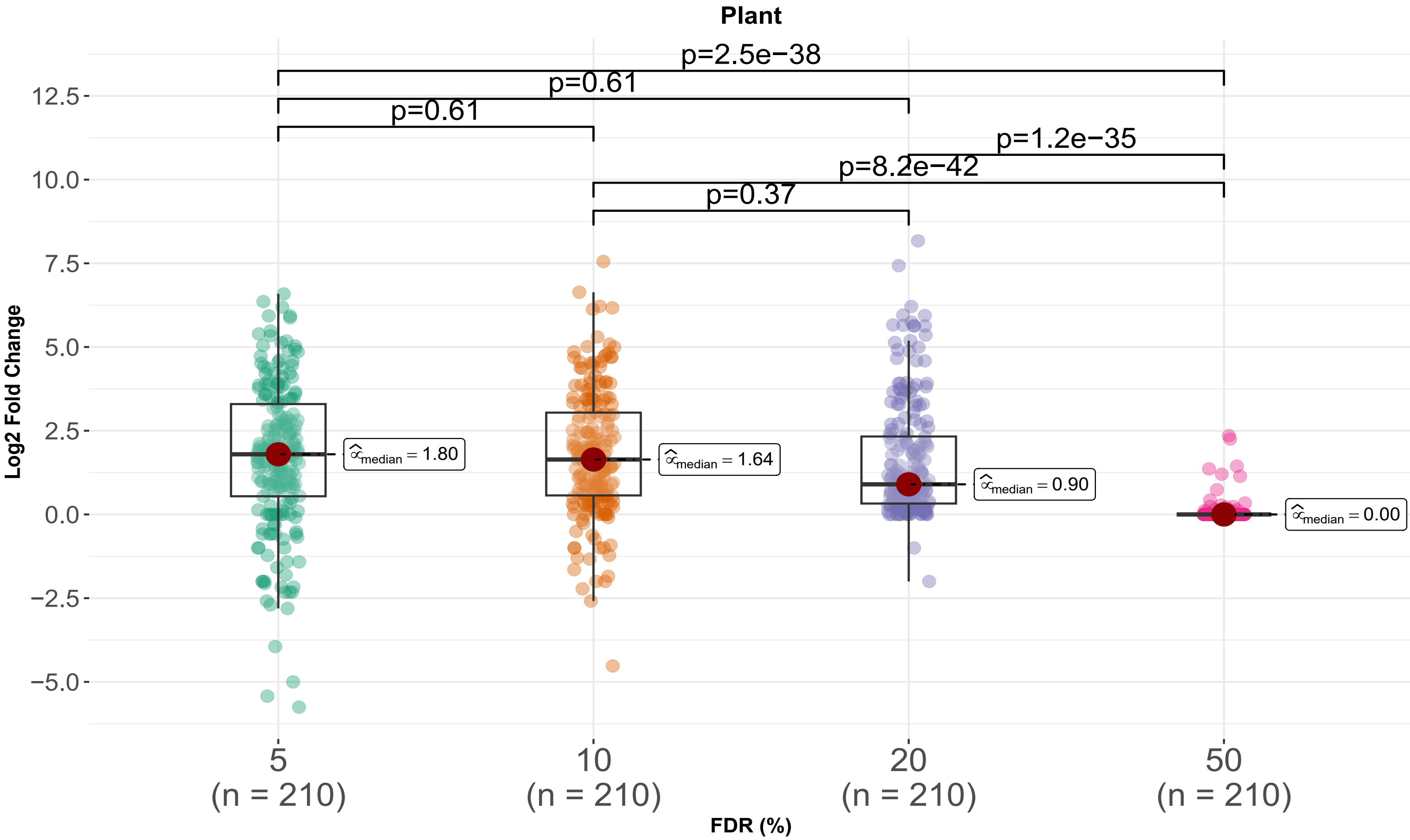


***Supplementary Fig. 15: Distribution of LFC in plant testing datasets across FDR thresholds***

Box plot showing distribution of Log2 fold changes of the number of annotations of METASPACE-ML relative to rule-based approach across all animal testing datasets, for different FDR thresholds. A dot corresponds to a test dataset. Exact p-values from a two-tailed Wilcoxon rank-sum test in (B) and (C) are shown above each comparison. Boxplots' bottom and top edges represent the 25th and 75th percentiles, with the median (50th percentile) line inside the box. Whiskers extend to the minimum and maximum values within 1.5 times the interquartile range from the quartiles; the minimum and maximum values are represented by the extent of the jittered data points.

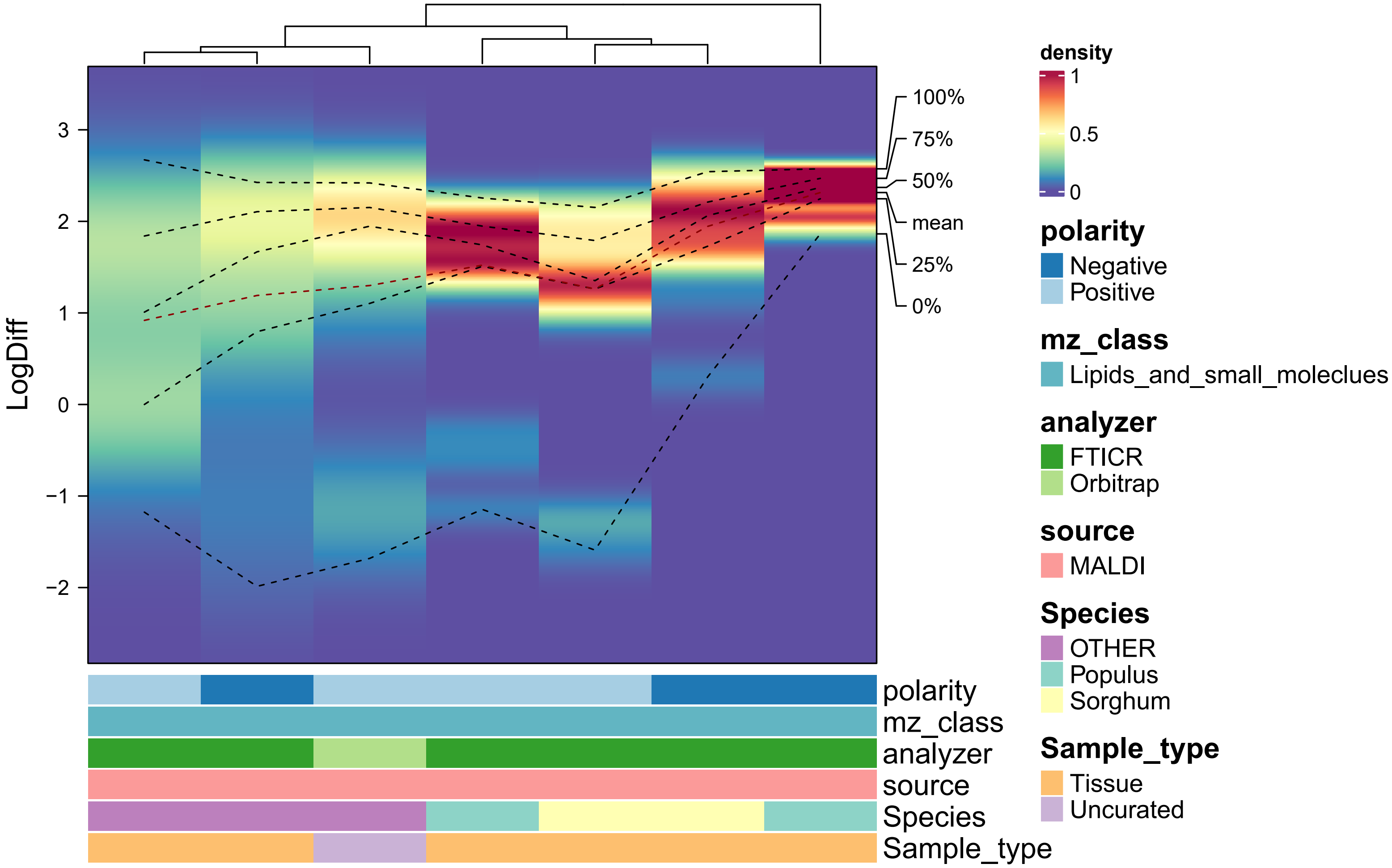


***Supplementary Fig. 16: Context-specific distribution of MAP scores for plant testing datasets***

Density heatmap showing the distribution of Log10 difference scores across datasets for each context in plant-based testing datasets. Each column represents a context which is described by its constituent metadata as 1D annotation bars colored by the classes in each metadata variable. The y-axis shows the MAP scores and the color gradient represents their density. Columns are hierarchically clustered using a distance metric based on the Kolmogorov-Smirnov statistic. Dotted lines in each column represent different quantiles of the MAP scores in addition to the mean.

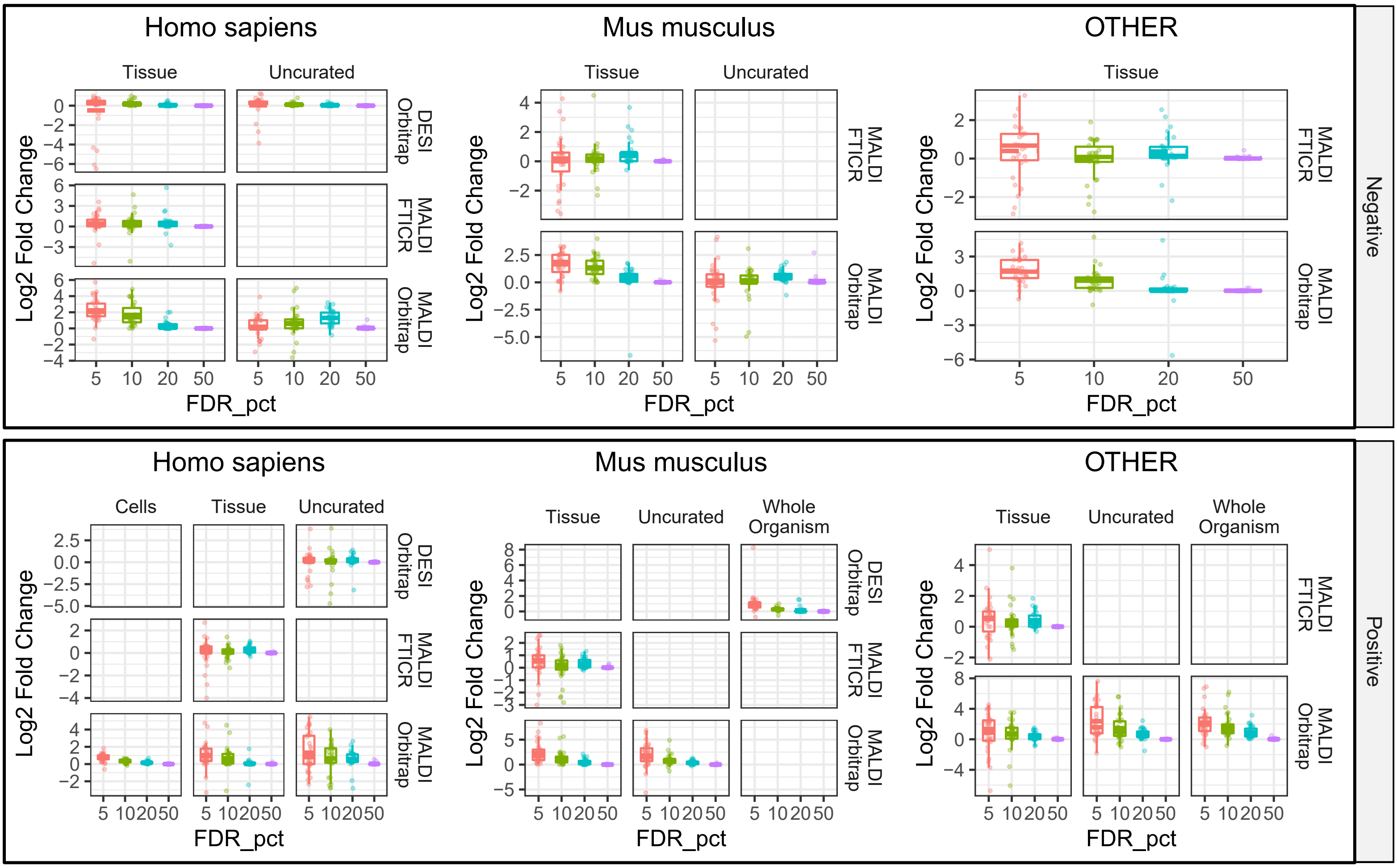


***Supplementary Fig. 17: Context-specific LFC scores for animal testing datasets across FDR thresholds***

Each box plot shows distribution of Log2 fold changes of the number of annotations of METASPACE-ML relative to rule-based approach across 30 datasets for each context across different FDR thresholds. Upper and lower panels represent negative and positive polarity, respectively. Each panel is divided into 3 sub panels based on the organism and each subpanel is a grid where columns and rows represent sample type and ionization source + analyzer, respectively. Boxplots' bottom and top edges represent the 25th and 75th percentiles, with the median (50th percentile) line inside the box. Whiskers extend to the minimum and maximum values within 1.5 times the interquartile range from the quartiles; the minimum and maximum values are represented by the extent of the jittered data points.

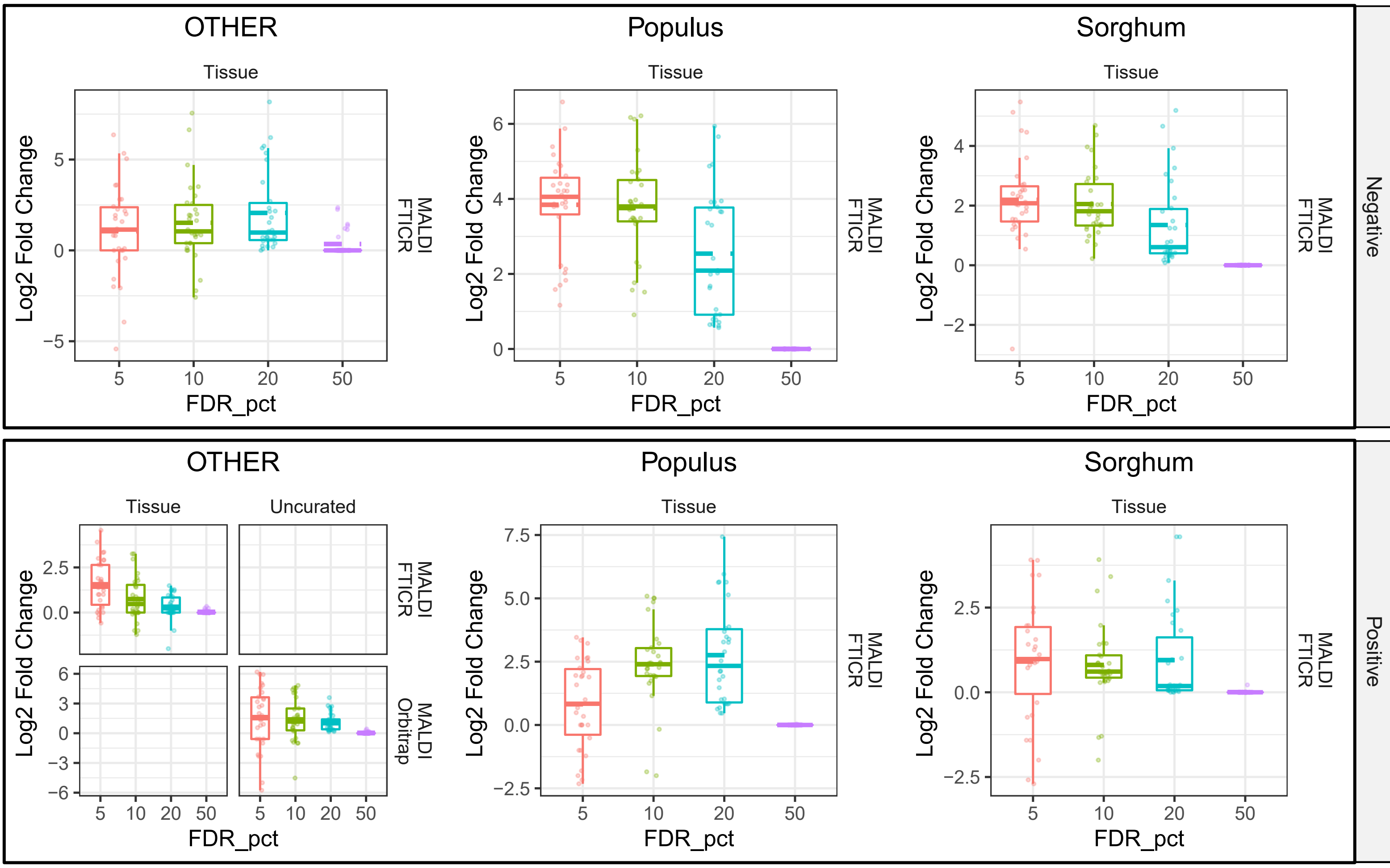


***Supplementary Fig. 18: Context-specific LFC scores for plant testing datasets across FDR thresholds***

Each box plot shows distribution of Log2 fold changes of the number of annotations of METASPACE-ML relative to rule-based approach across 30 datasets for each context across different FDR thresholds. Upper and lower panels represent negative and positive polarity, respectively. Each panel is divided into 3 sub panels based on the organism and each subpanel is a grid where columns and rows represent sample type and ionization source + analyzer, respectively. Boxplots' bottom and top edges represent the 25th and 75th percentiles, with the median (50th percentile) line inside the box. Whiskers extend to the minimum and maximum values within 1.5 times the interquartile range from the quartiles; the minimum and maximum values are represented by the extent of the jittered data points.

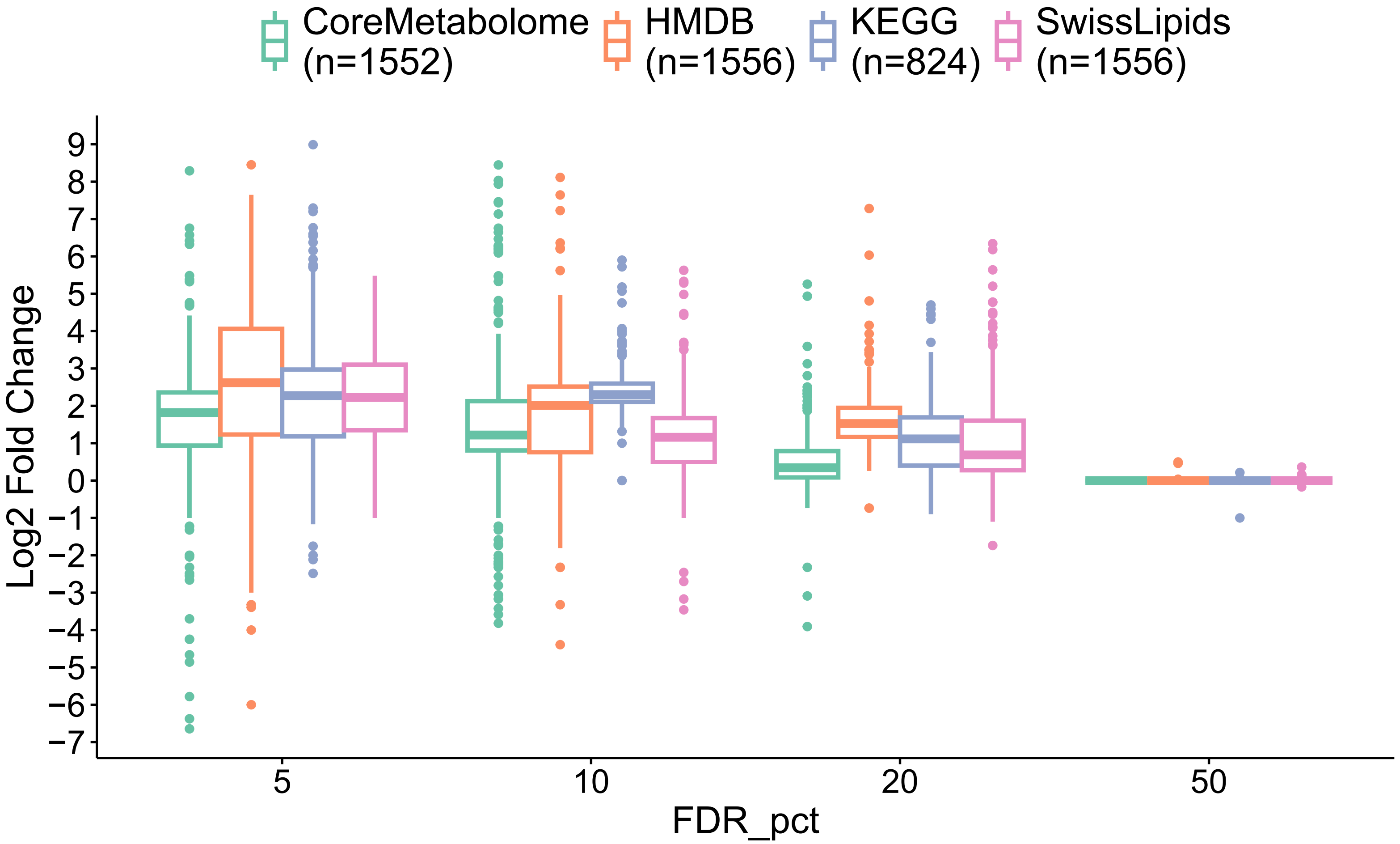


***Supplementary Fig. 19: Distribution of LFC scores across FDR thresholds for each annotation database***

Boxplot showing the log2 fold change (LFC) between the number of annotations captured by METASPACE-ML compared to MSM-based approach across different FDR % thresholds for each annotation database. Boxplot color corresponds to annotation database. Number of dataset groups per database are displayed below legend labels. Boxplots' bottom and top edges represent the 25th and 75th percentiles, with the median (50th percentile) line inside the box. Whiskers extend to the minimum and maximum values within 1.5 times the interquartile range from the quartiles; the minimum and maximum values are represented by the extent of the jittered data points.

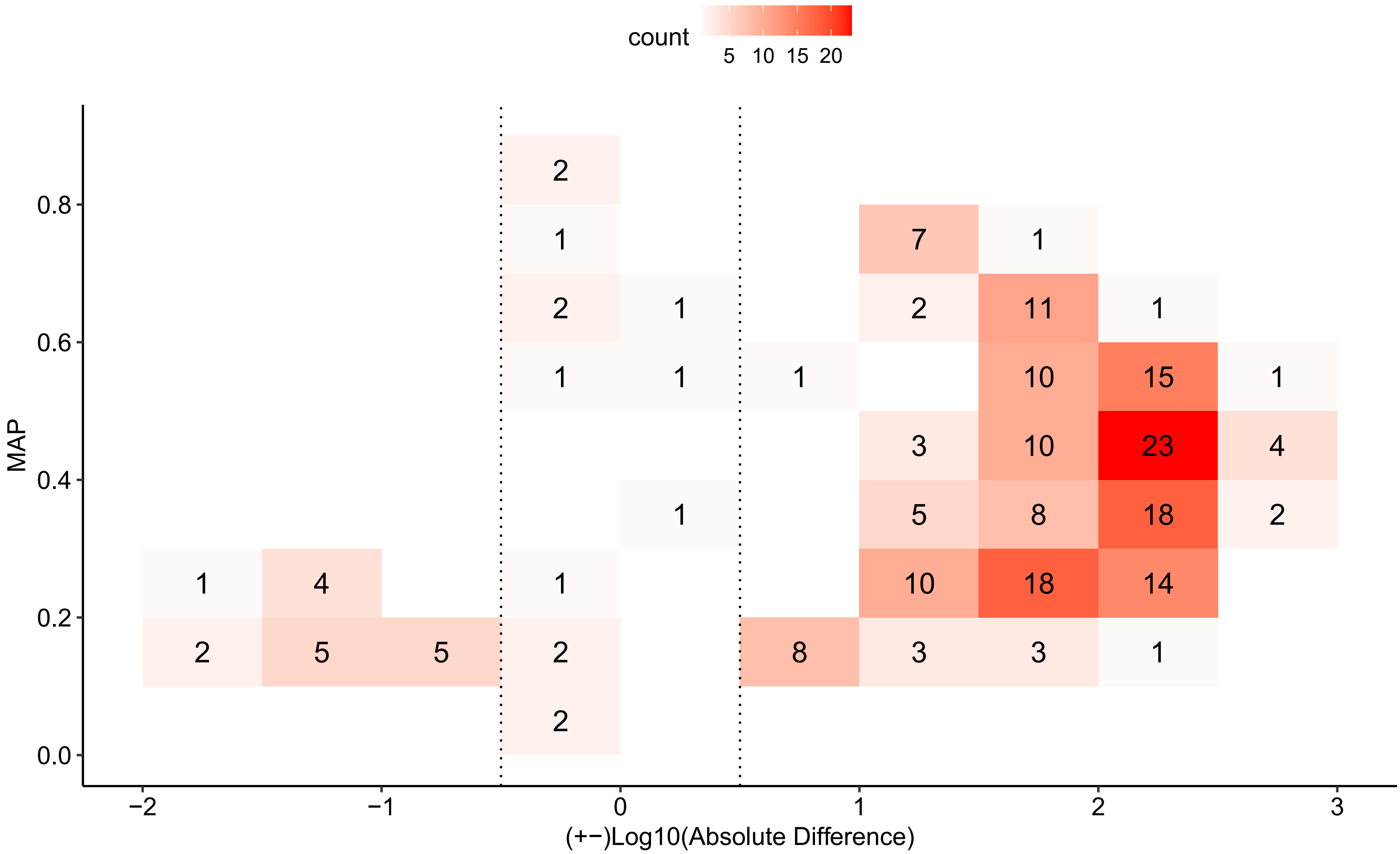


***Supplementary Fig. 20: Relationship between Log10 absolute difference score and MAP scores in plant testing datasets***

Relationship between Log10 absolute difference score and MAP scores depicted by a 2D rectangular binned plot. Each bin has a width of 0.5 and a length of 0.1 corresponding to x-axis and y-axis breaks, respectively. The color gradient corresponds to the number of plant testing datasets within each bin and the count is also displayed per bin. Bins within the dotted lines represent datasets with <= 0 Log10 absolute difference scores.

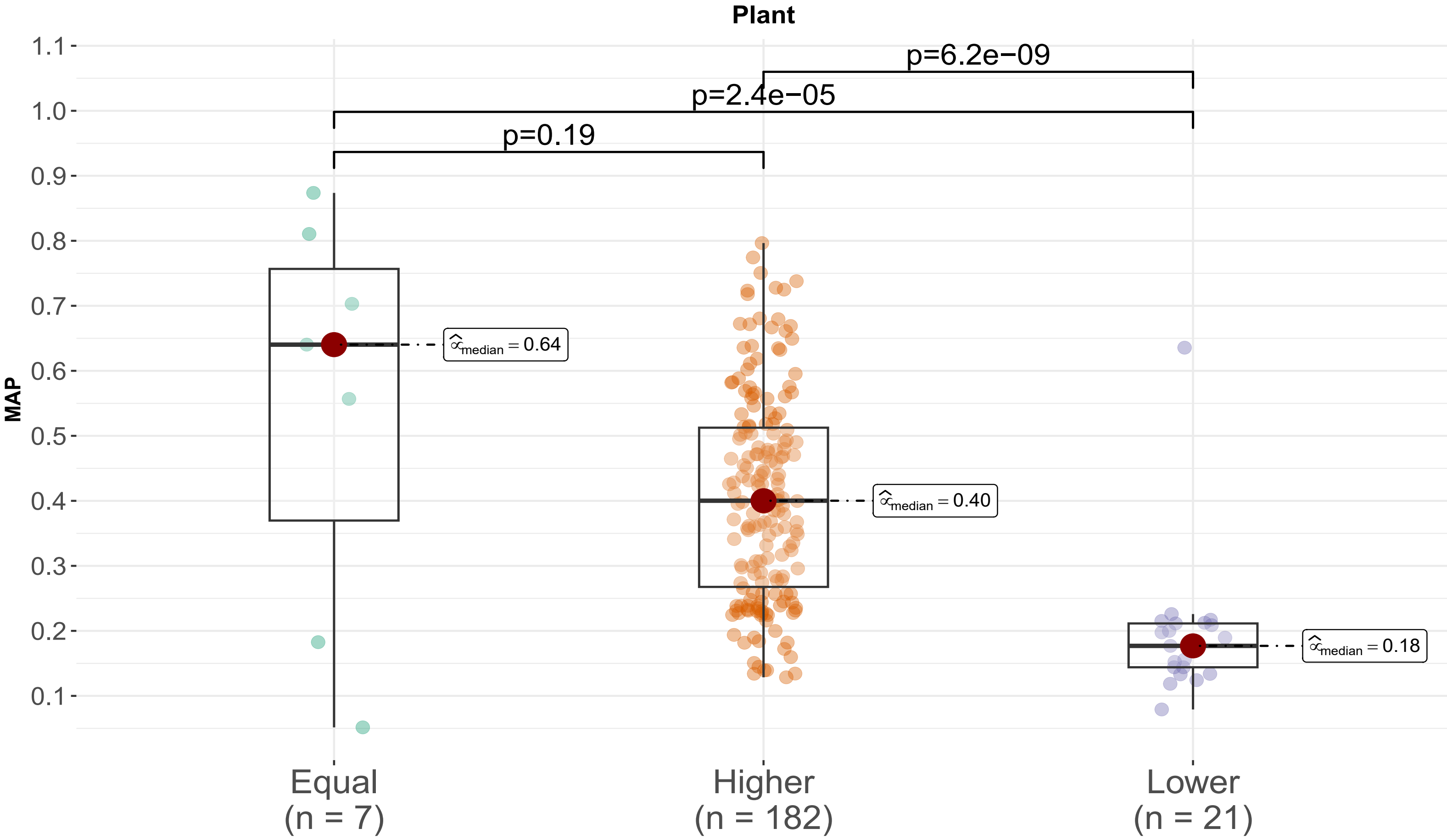


***Supplementary Fig. 21: Comparison of MAP scores within plant testing datasets according to annotation coverage***

Box plot showing distribution of MAP scores across animal testing datasets and compared across different groups based on whether METASPACE-ML had equal, higher or lower annotations compared to rule-based approach. Exact p-values from a two-tailed Wilcoxon rank-sum test in (B) and (C) are shown above each comparison. Boxplots’ bottom and top edges represent the 25th and 75th percentiles, with the median (50th percentile) line inside the box. Whiskers extend to the minimum and maximum values within 1.5 times the interquartile range from the quartiles; the minimum and maximum values are represented by the extent of the jittered data points.

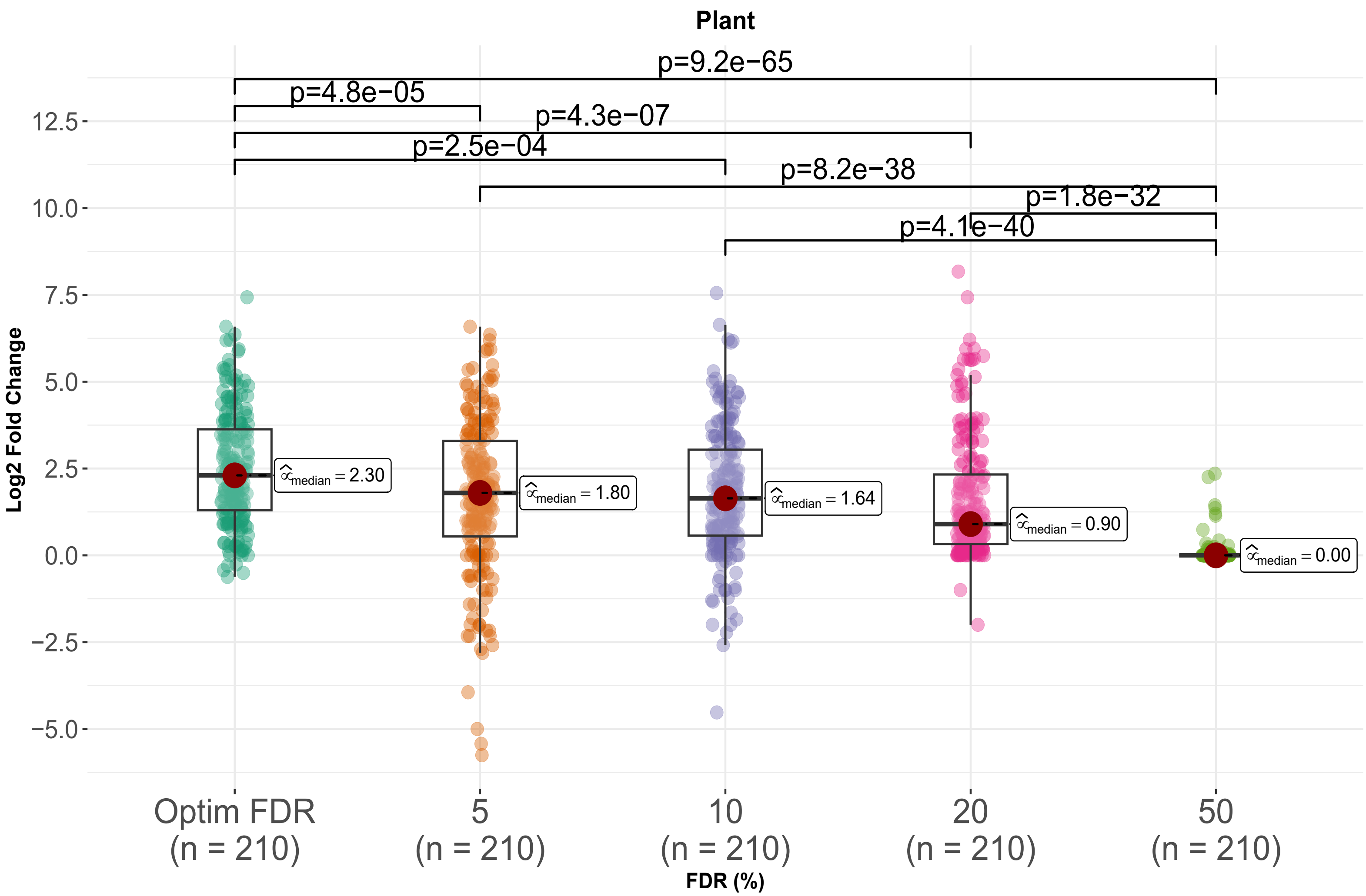


***Supplementary Fig. 22: Distribution of LFC in plant testing datasets across FDR thresholds including optimized FDR***

Box plot showing distribution of Log2 fold changes of the number of annotations of METASPACE-ML relative to rule-based approach across all plant testing datasets, for different FDR thresholds. A dot corresponds to a test dataset. Exact p-values from a two-tailed Wilcoxon rank-sum test in (B) and (C) are shown above each comparison. Boxplots' bottom and top edges represent the 25th and 75th percentiles, with the median (50th percentile) line inside the box. Whiskers extend to the minimum and maximum values within 1.5 times the interquartile range from the quartiles; the minimum and maximum values are represented by the extent of the jittered data points.

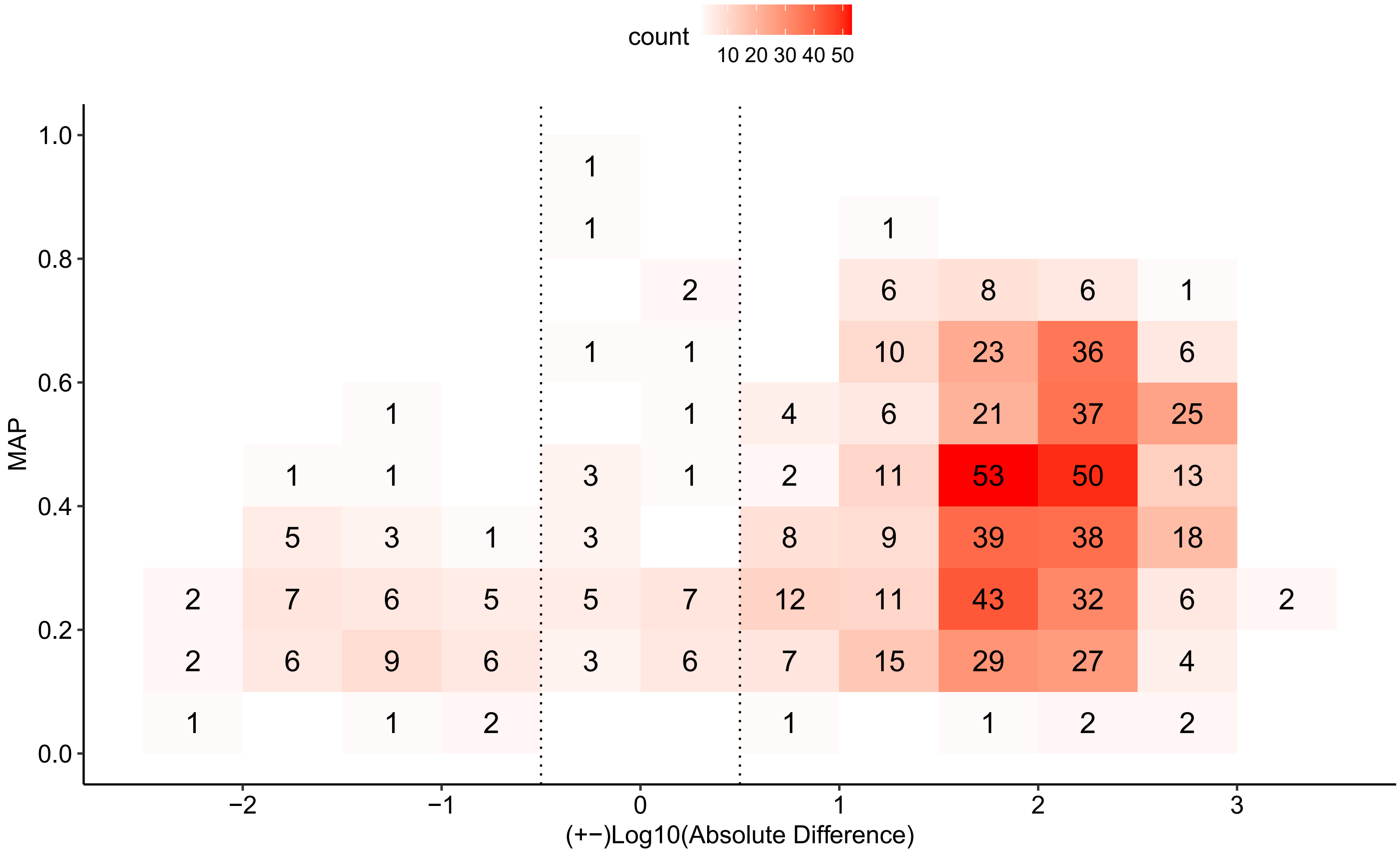


***Supplementary Fig. 23: Relationship between Log10 absolute difference score and MAP scores in animal testing datasets after FDR threshold optimization.***

Relationship between Log10 absolute difference score and MAP scores depicted by a 2D rectangular binned plot. Each bin has a width of 0.5 and a length of 0.1 corresponding to x-axis and y-axis breaks, respectively. The color gradient corresponds to the number of animal testing datasets within each bin and the count is also displayed per bin. Bins within the dotted lines represent datasets with <= 0 Log10 absolute difference scores.

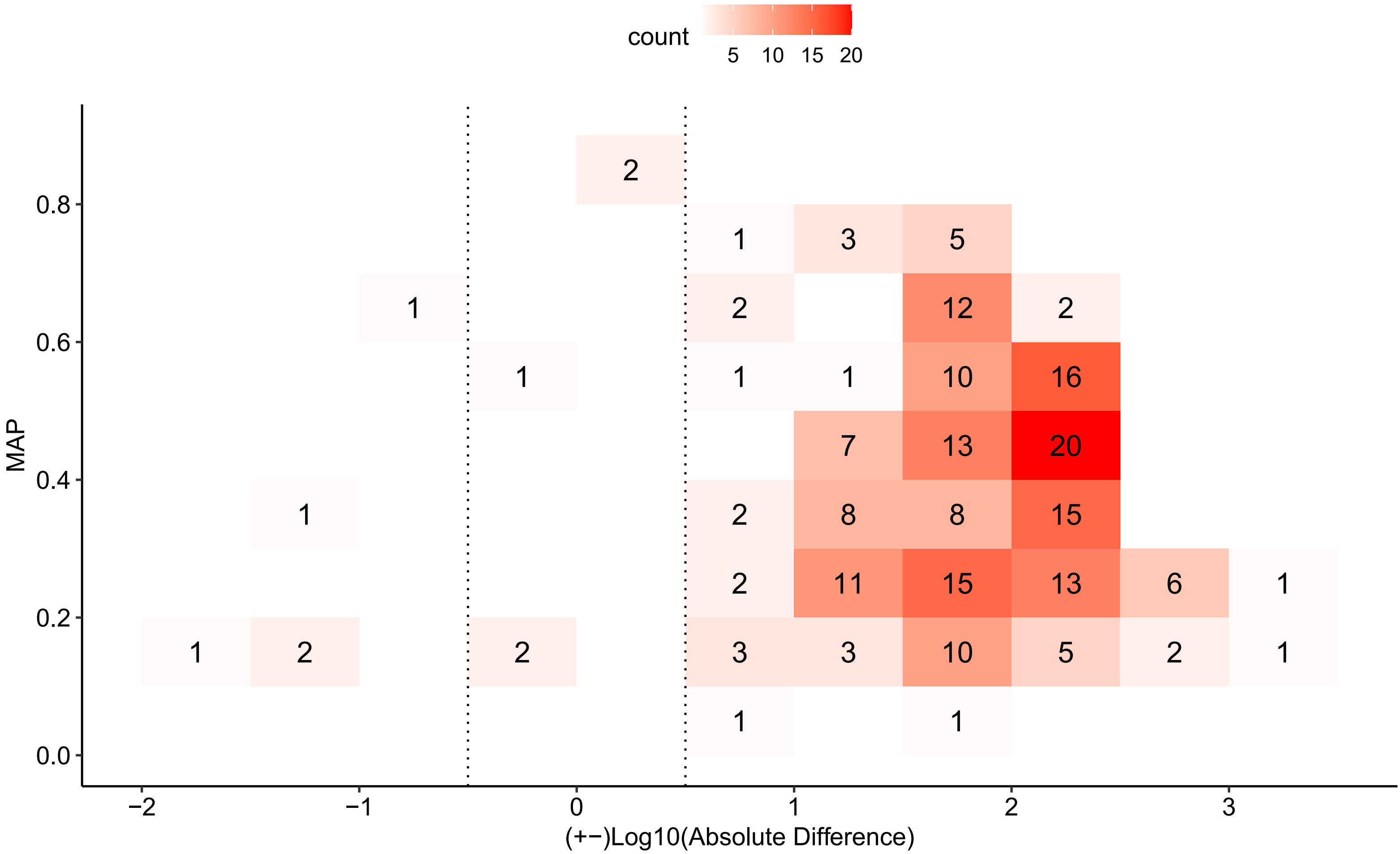


***Supplementary Fig. 24: Relationship between Log10 absolute difference score and MAP scores in plant testing datasets after FDR threshold optimization.***

Relationship between Log10 absolute difference score and MAP scores depicted by a 2D rectangular binned plot. Each bin has a width of 0.5 and a length of 0.1 corresponding to x-axis and y-axis breaks, respectively. The color gradient corresponds to the number of plant testing datasets within each bin and the count is also displayed per bin. Bins within the dotted lines represent datasets with <= 0 Log10 absolute difference scores.

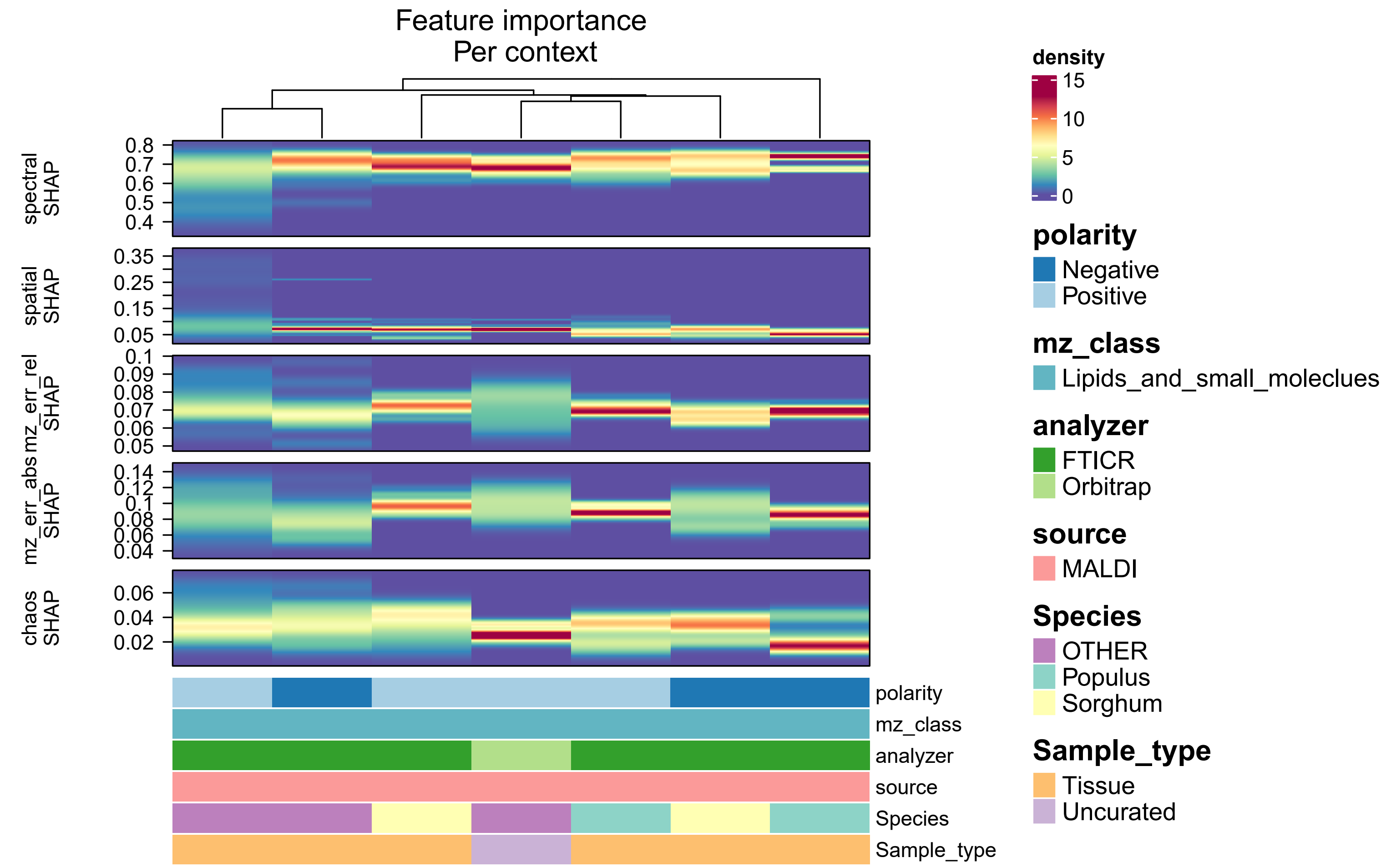


***Supplementary Fig. 25: Feature importance in plant testing datasets***

Density heatmap showing the distribution of SHAP impact contributions scores across datasets for each context in plant-based testing datasets and faceted by each of the 5 features. Each column represents a context which is described by its constituent metadata as 1D annotation bars colored by the classes in each metadata variable. The y-axis shows the SHAP impact contributions scores and the color gradient represents their density. Columns are hierarchically clustered using a distance metric based on the Kolmogorov-Smirnov statistic.

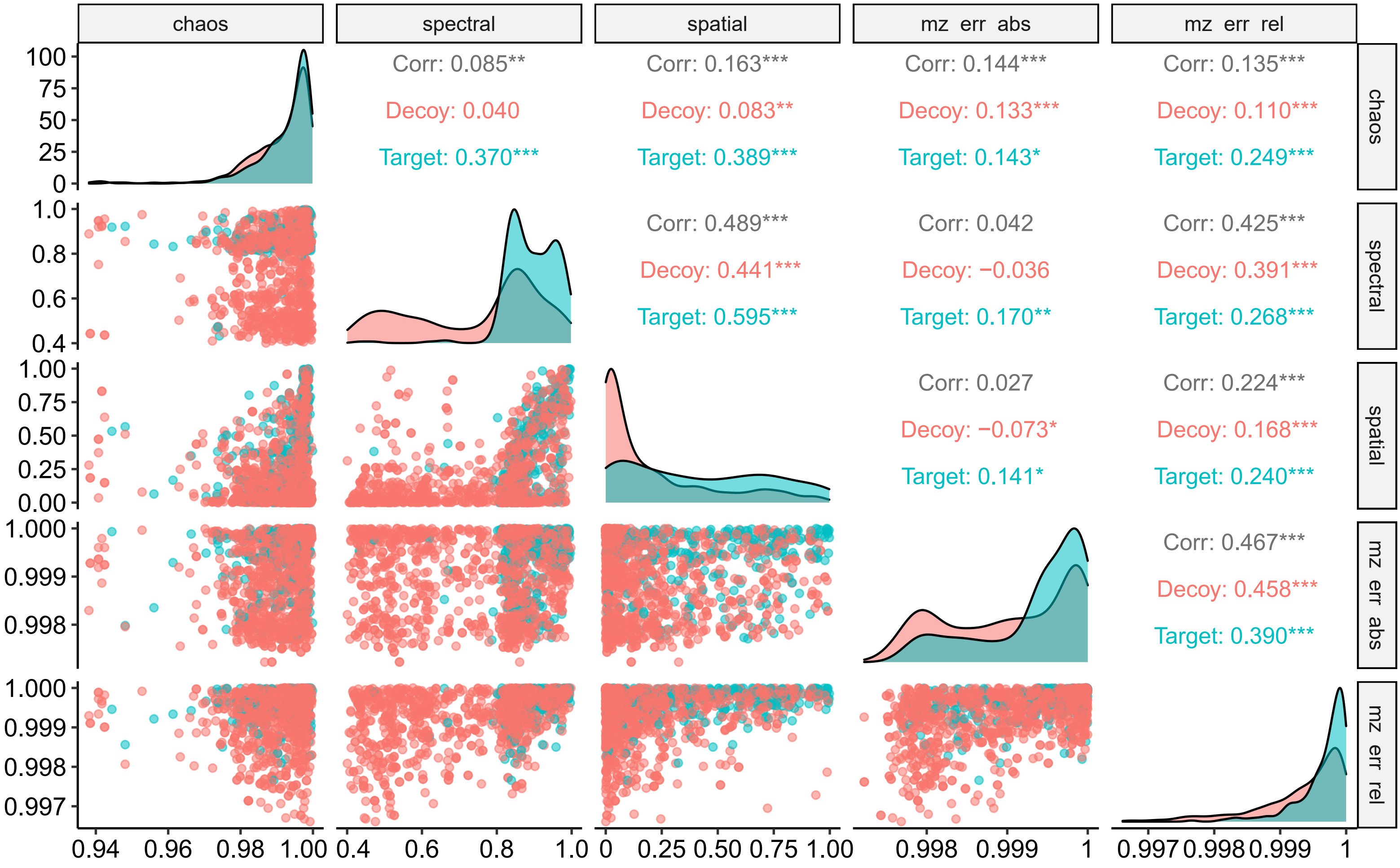


***Supplementary Fig. 26: Pairwise relationships between model features for a mouse brain dataset***

Scatterplot matrix plotting pairwise feature scores for target and decoy ions for one of the brain datasets used for bulk validation (https://metaspace2020.eu/dataset/2016-09-22_11h16m09s). Columns and rows denote the 5 features used to train the METASPACE-ML model. The lower and upper triangles show the scatter plots and Pearson correlation coefficients with significance asterisks, respectively. Each dot represents an ion and is colored by whether it's a target or decoy as well as the correlation coefficients. A density plot is drawn along the diagonal for each feature and is also grouped by ion type (target vs decoy). Asterisks are added to denote significance level (*** < 0.001, ** < 0.01, * < 0.05).

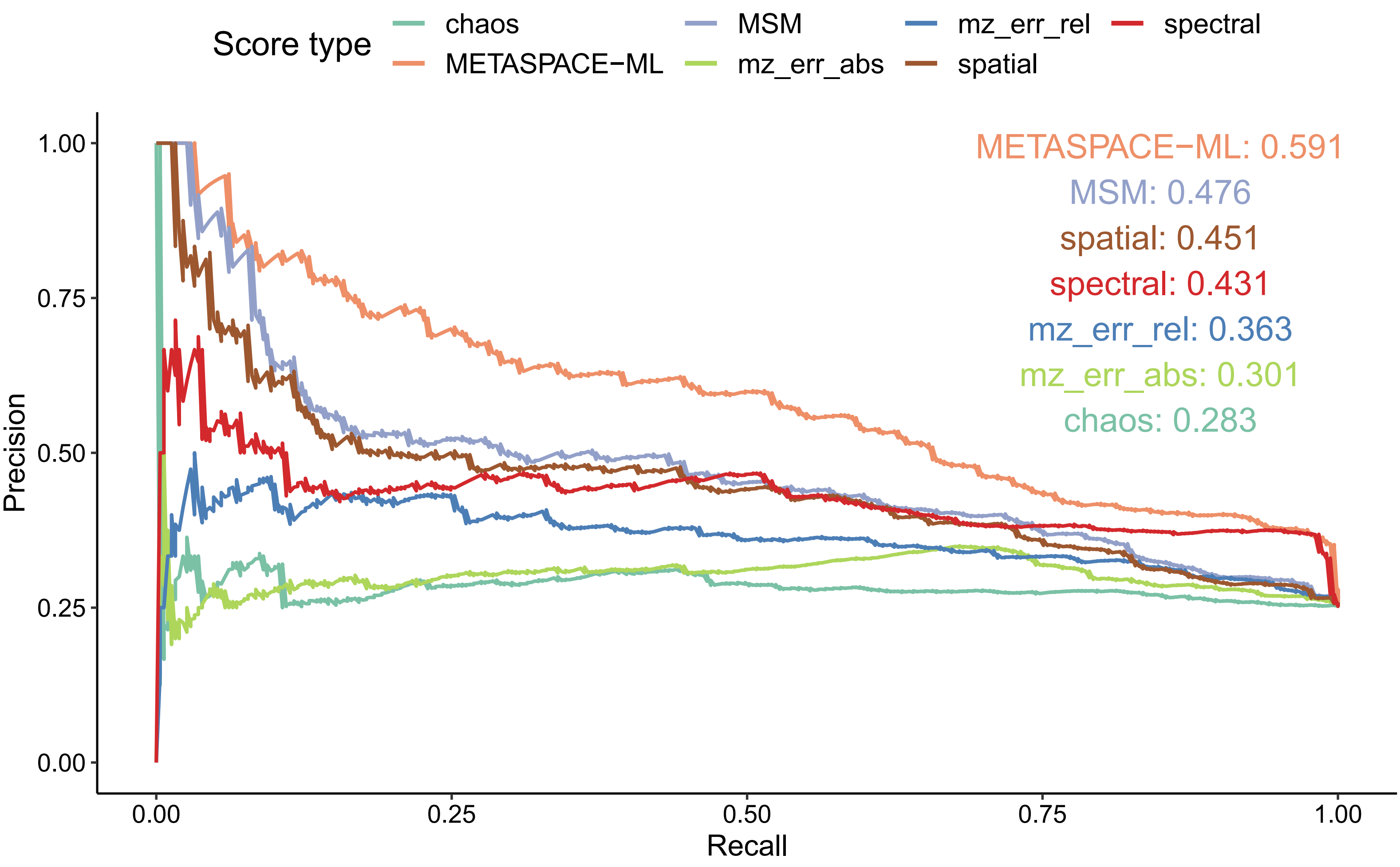


***Supplementary Fig. 27: Precision-Recall curves for a mouse brain dataset***

Precision-Recall curves for the same dataset as in (Supplementary Fig. 26) showing precision and recall using METASPACE-ML and MSM scores in addition to the 5 constituent features. Curve color corresponds to score type.

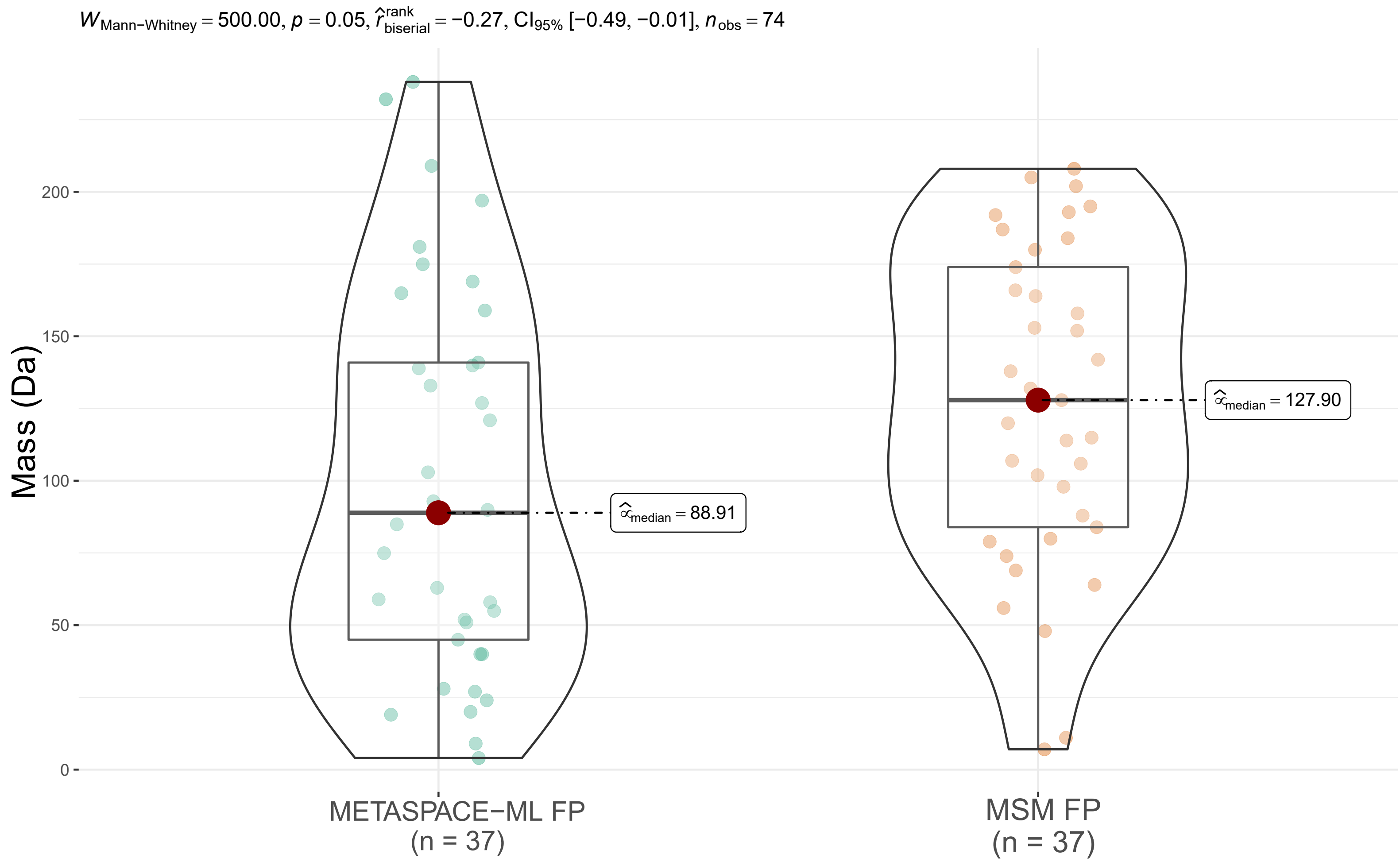


***Supplementary Fig. 28: Comparison of decoys' exact masses between METASPACE-ML and MSM false positives.***

For each decoy adduct associated with a decoy ion having FDR < 10%, false positives were classified into METASPACE-ML FP if the mean Log10 absolute difference scores across animal testing datasets is < 0 and MSM FP otherwise. The box-violin plot above shows distribution of those adduct exact masses for both groups. The p-value and Mann-Whitney statistics are displayed as a subtitle above the plot and the number of adducts per group is displayed in parentheses below the x-axis labels. Boxplots' bottom and top edges represent the 25th and 75th percentiles, with the median (50th percentile) line inside the box. Whiskers extend to the minimum and maximum values within 1.5 times the interquartile range from the quartiles; the minimum and maximum values are represented by the extent of the jittered data points.

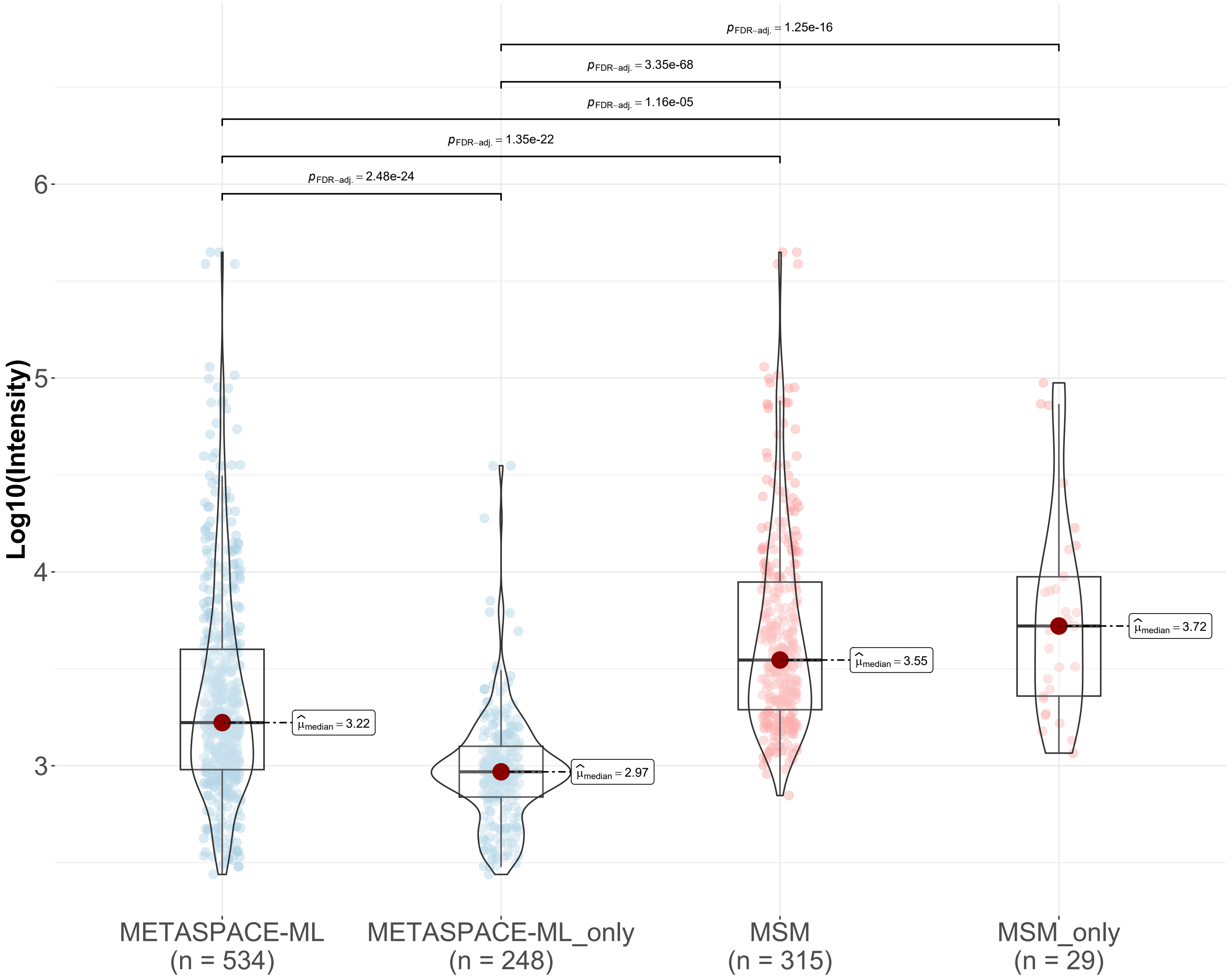


***Supplementary Fig. 29: Distribution of ion intensities in a mouse brain dataset.***

Box-violin plot showing the distribution of ion intensities in the dataset (https://metaspace2020.eu/dataset/2019-08-19_11h28m42s). Y-axis corresponds to log10 total intensity of each ion, x-axis represents the respective groups of ions which also includes the number of ions in each group enclosed in parentheses. Significance results of pairwise comparisons between groups are displayed above each respective pair of groups only if the result is significant (p-value < 0.05) according to Kruskal-Wallis test. The median of each distribution is also indicated next to each boxplot. Boxplots' bottom and top edges represent the 25th and 75th percentiles, with the median (50th percentile) line inside the box. Whiskers extend to the minimum and maximum values within 1.5 times the interquartile range from the quartiles; the minimum and maximum values are represented by the extent of the jittered data points.

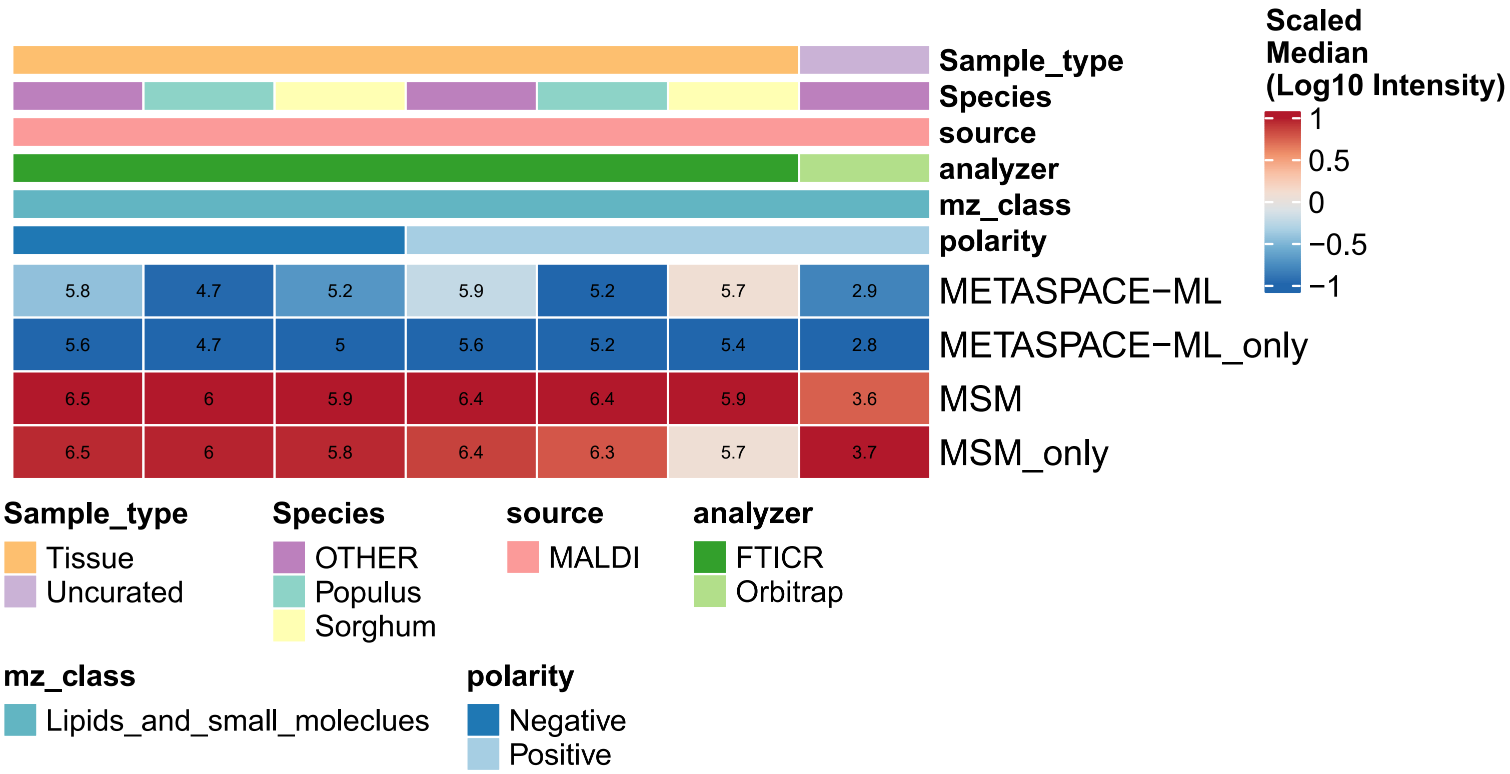


***Supplementary Fig. 30: Context-specific intensity comparison between different approaches in plant testing datasets***

Heatmap showing the median log10 intensity per context across plant testing datasets. Each column represents a context which is described by its constituent metadata as 1D annotation bars colored by the classes in each metadata variable and rows representing different model approaches where the suffix “only” indicate that annotations were exclusively captured by a given approach and not the other. Color gradient and labels correspond to median log10 intensity.

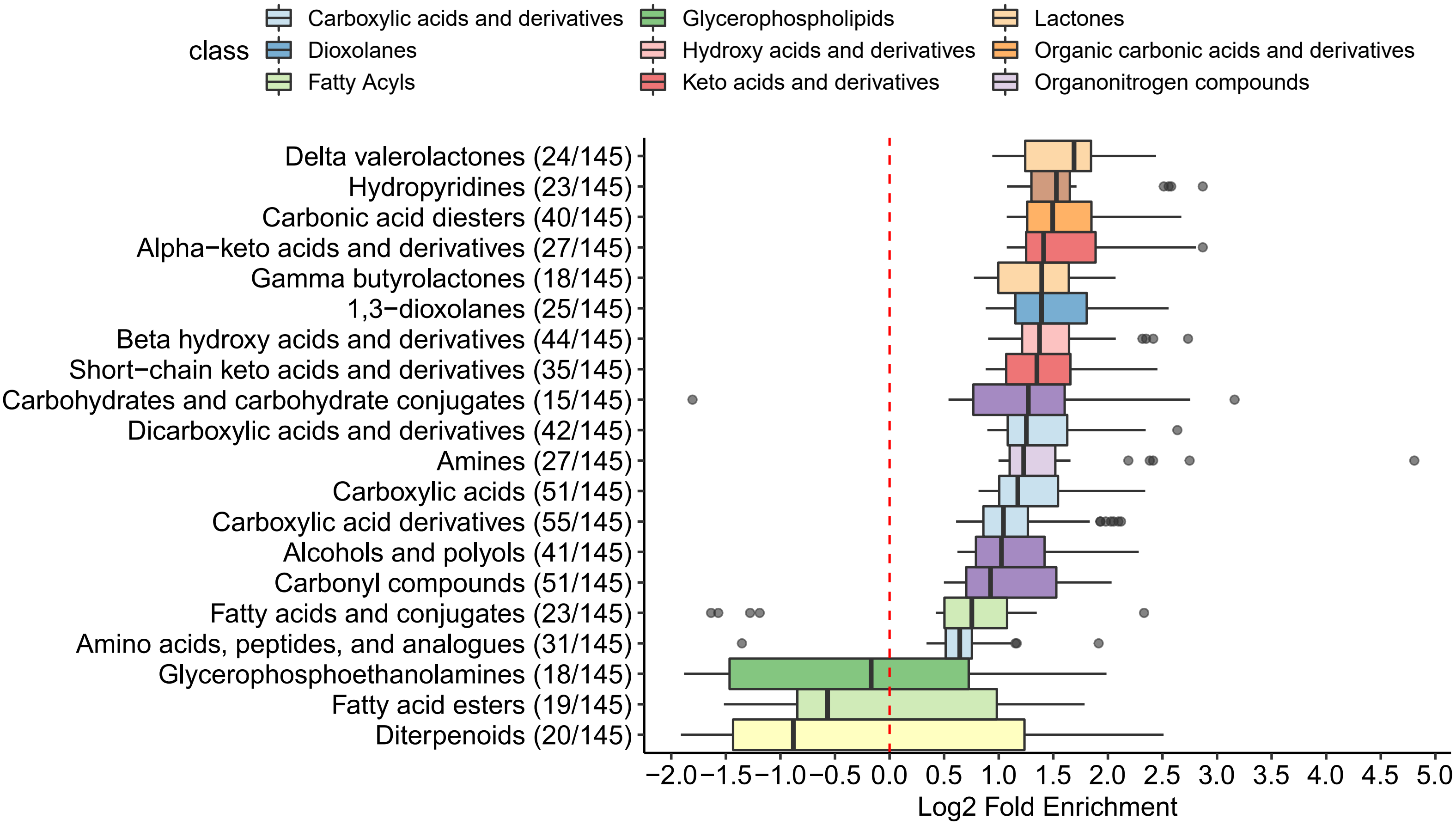


***Supplementary Fig. 31: Overall metabolic class enrichment in plant testing datasets***

Overrepresentation analysis was performed using Fisher-exact test for each dataset to check whether there are specific metabolite/lipid classes enriched in the ions that are only captured by METASPACE-ML at 10% FDR. Log2 fold enrichment (See Methods) is plotted on the x-axis and HMDB metabolite subclasses are plotted on the y-axis with the number of datasets represented in each boxplot enclosed in parentheses compared to the number of datasets with at least one significant term ($p$-value $< 0.05$ and with a minimum intersections size of 3 ions). Boxplots are colored according to the parent class where each subclass belongs to. Only terms having significant enrichment ($p$-value $< 0.05$) in at least 10% of all significant datasets are shown. Boxplots' bottom and top edges represent the 25th and 75th percentiles, with the median (50th percentile) line inside the box. Whiskers extend to the minimum and maximum values within 1.5 times the interquartile range from the quartiles; the minimum and maximum values are represented by the extent of the jittered data points.

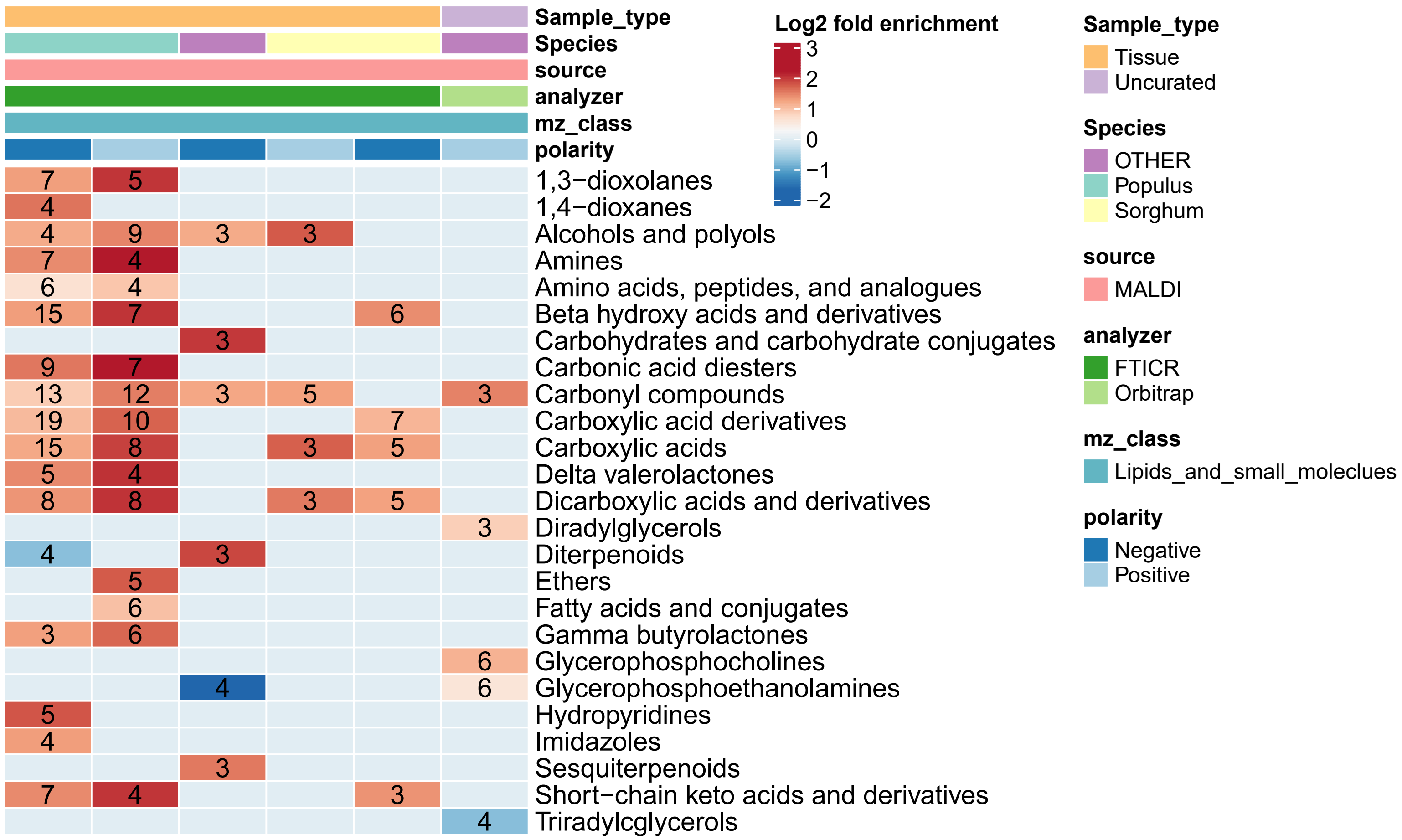


***Supplementary Fig. 32: Context-specific metabolic class enrichment in plant testing datasets***

Heatmap showing the overrepresentation analysis results per context in plant testing datasets. Each column represents a context which is described by its constituent metadata as 1D annotation bars colored by the classes in each metadata variable and rows representing metabolite classes significantly enriched in a given context. Color gradient corresponds to Log2 fold enrichment and labels corresponding to number of datasets per context where a given class is enriched.

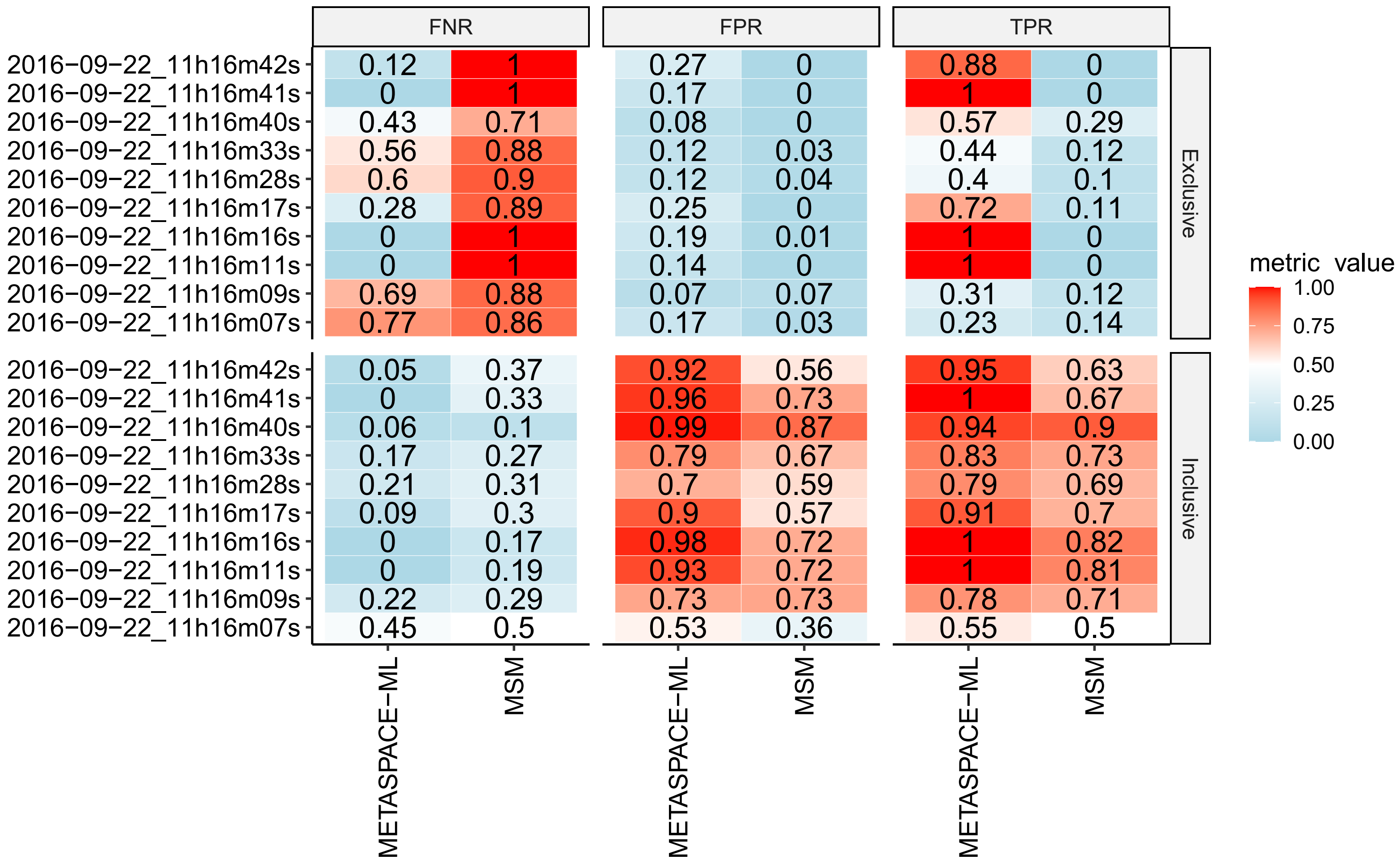


***Supplementary Fig. 33: Diagnostic metrics for bulk validation datasets***

Tiled grid showing 3 diagnostic metrics : true positive rate (TPR), false positive rate (FPR) and false negative rate (FNR) displayed as columns for each of the 10 brain datasets used for validation displayed as rows. X-axis shows both the METASPACE-ML and MSM approach and color gradient corresponding to metrics' scores which have a scale of 0-1.
